# Dopaminergic neuromodulation of spike timing dependent plasticity in mature adult rodent and human cortical neurons

**DOI:** 10.1101/2020.05.15.098400

**Authors:** Emma Louise Louth, Rasmus Langelund Jørgensen, Anders Rosendal Korshøj, Jens Christian Hedemann Sørensen, Marco Capogna

## Abstract

Synapses in the cerebral cortex constantly change and this dynamic property regulated by the action of neuromodulators such as dopamine (DA), is essential for reward learning and memory. DA modulates spike-timing-dependent plasticity (STDP), a cellular model of learning and memory, in juvenile rodent cortical neurons. However, it is unknown whether this neuromodulation also occurs at excitatory synapses of cortical neurons in mature adult mice or in humans. Cortical layer V pyramidal neurons were recorded with whole cell patch clamp electrophysiology and an extracellular stimulating electrode was used to induce STDP. DA was either bath-applied or optogenetically released in slices from mice. Classical STDP induction protocols triggered non-Hebbian excitatory synaptic depression in the mouse or no plasticity at human cortical synapses. DA reverted long term synaptic depression to baseline in mouse or elicited long term synaptic potentiation in human cortical synapses. Furthermore, when DA was applied during a STDP protocol it depressed presynaptic inhibition in the mouse but not in the human cortex. Thus, DA modulates excitatory synaptic plasticity differently in human *versus* mouse cortex. The data strengthens the importance of DA in gating cognition in humans, and may inform on therapeutic interventions to recover brain function from diseases.

## Introduction

Humans and other mammalians are characterized by their ability to produce goal-directed and intelligent behaviors beyond simple stimulus–response associations. It is believed that the neuromodulator dopamine (DA) plays a key role in gating cortical operations underlying cognitive functions, such as working memory ^1^, attention ^2^, and flexible behavior ^3^. Furthermore, evidence suggests that behaviorally-relevant sensory information and contextual information are gated by DA to maintain relevant information in working memory and relay choice signals ^4^.

DA is synthesized by specific neurons in the midbrain that send their widespread projections to several brain regions including the cerebral cortex ^5^. The gating elicited by DA, released by midbrain axonal varicosities and terminal endings within the cerebral cortex, is assumed to represent a key molecular substrate underlying cognitive performance including stimulus selection, working memory, rule switching and decision making ^6^.

Many theories have been proposed to account for by DA circuit mechanisms underlying cortical-mediated executive control ^4^. One of the most successful is the reward prediction error theory and its experimental demonstration in DA neurons ^7^

A phenomenon that could represent a landmark cellular substrate of cognitive functions is represented by the DA modulation of spike timing dependent plasticity (STDP). STDP is a form of synaptic plasticity triggered by repeated pairings of single presynaptic and postsynaptic spikes ^8–11^. It depends on the order and millisecond-precision timing of spikes: multiple pre- before-post spike pairings often evoke timing-dependent long-term potentiation (t-LTP), whereas post-before-pre pairings often evoke timing-dependent long-term depression (t-LTD). STDP is a remarkable example of Hebbian plasticity ^12^, since synaptic inputs that promote postsynaptic firing are strengthened. However, non-Hebbian STDP has also been observed, that is multiple pre before-post spike pairings elicit t-LTD ^13^.

It is well established that DA has an important modulatory role on STDP, as it broadens the time window for detecting coincident spiking in the pre- and postsynaptic neurons and in this way boosts the induction of t-LTP in rodent neocortical neurons ^14–16^. Furthermore, DA modulates the polarity of STDP promoting t-LTP at excitatory synapses of rodent prefrontal cortex (PFC) ^16^ and hippocampus ^14,17–20^.

However, there are two main drawbacks with the DA modulation of STDP model. First, STDP is often considered to represent a cellular substrate for cognitive operations performed by adult mammalians. Yet, most of the data have been obtained from developmental and juvenile rodents (usually 2-3 weeks old) including those studying DA modulation of STDP ^21^. Second, researchers wish to use STDP to model synaptic plasticity occurring at human cortical neurons ^22^, since this form of synaptic plasticity can be demonstrated at excitatory synapses of human cortex ^23^. However, it is still unknown whether DA modulates STDP at human neocortical synapses.

We aimed to make progress on these two issues by testing the DA modulation of STDP from neurosurgically-resected adult human neocortical slices and by comparing it to the DA modulation of STDP in mature adult mouse cortical synapses.

## Results

### Excitatory synapses show t-LTD in mature adult mice

In order to characterize STDP in mature adult mice, we tested different STDP timings (Δt = −30 ms, −10 ms, +10 ms and +30 ms) in 60-80 day old mice (Fig 1A). We patched layer V pyramidal neurons in whole-cell configuration and placed a stimulating electrode approximately 100-150 μm from the soma, nearby the apical dendrite, without pharmacological blockade of synaptic inhibition to mimic physiological conditions. We first recorded a 10 minute baseline of evoked excitatory postsynaptic potentials (EPSPs), followed by an STDP induction protocol consisting of 75 EPSPs to action potential (AP) pairings. We then monitored evoked EPSPs for another 40 minutes (Fig 1B). We did not observe significant effects induced by Δt 30 ms (Fig 1 A, −30 ms: 85.9 ± 10.4% vs 100 %, n = 6, *p* = 0.2, paired *t*-test; +30 ms: 87.2 ± 10.1% vs 100 %, n = 6, *p* = 0.3). We also observed no change at Δt −10 ms (Fig 1 A, 105.8 ± 9.5% vs 100 %, n = 6, *p* = 0.6). In contrast, at Δt +10 ms neurons exhibited t-LTD where EPSPs had a decreased rising slope (Fig 1 A and B, 53.6 ± 8.4% vs 100 %, n = 8, *p* = 0.009) and peak amplitude (Figure Supplement 1). Interestingly, we found that at mature adult excitatory synapses, the plasticity outcome was dependent on the strength of postsynaptic depolarization, as reported for younger mice ^24^. Experimentally, we replaced a single spike Δt +10 ms induction with a 25 ms depolarization protocol to produce 3-4 APs as shown in Figure Supplement 2. We found that such burst protocol instead of producing t-LTD, as observed with a single spike protocol, resulted in no change in EPSP rising slope or peak amplitude (Figure Supplement 2).

**Fig 1.**
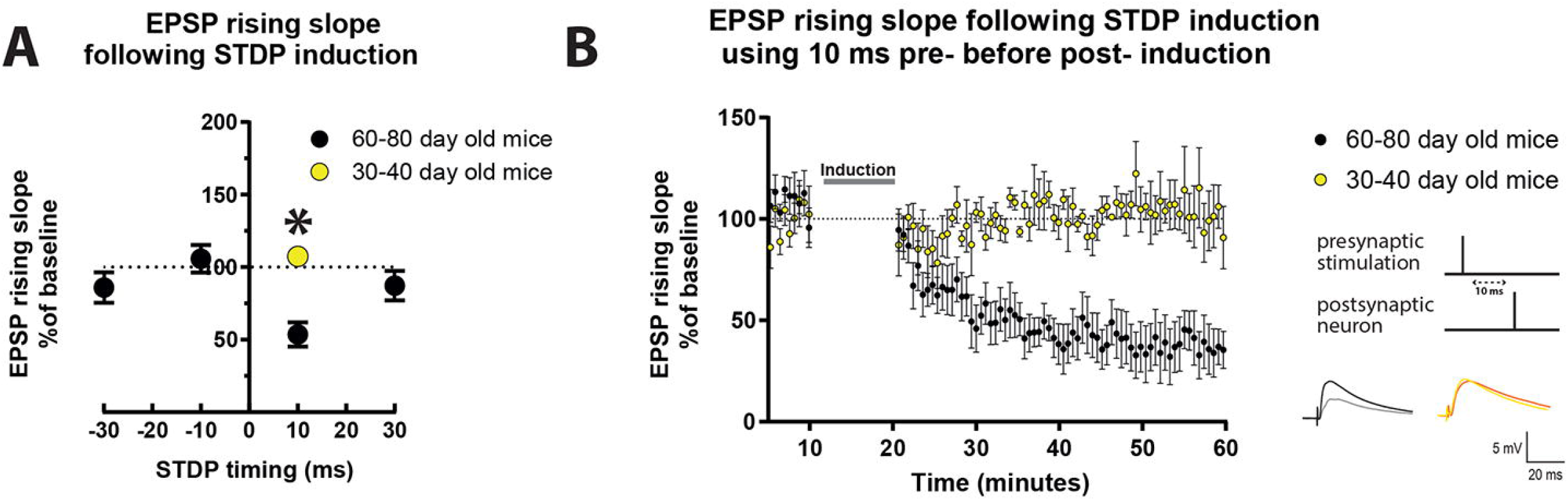
STDP induction at varying timings in layer 5 cortical pyramidal neurons of mature adult mice and comparison to adult mice. (A) STDP induction at EPSP-AP pairing timings (Δt) of −30 ms, −10 ms and +30 ms show no change in EPSP rising slope. At +10 ms neurons from mature adult mice (60-80 day old) exhibit t-LTD (*p* = 0.009 vs 100 %,) which was significantly different from adult mice (30-40 day old) (*p* = 0.001,). Neurons from adult mice exhibited a trend toward t-LTP (*p* = 0.08 vs 100 %). (B) The graph shows the time-course of the EPSP rising slope during the STDP experiments performed in both mature adult and adult mice. The grey bar in the graph illustrates the time of STDP induction. To the right, the STDP induction protocol timing is illustrated and below are example traces of EPSPs. The baseline trace is the darker trace, the traces following STDP induction are the lighter traces; each trace is the average of 80 traces from the same recording. All data are shown as mean ± SEM. Number of neurons recorded in mature adult mice, n = 8, and in adult mice n = 4.

Next, we compared the effect observed after the Δt +10 ms STDP protocol in mature adult mice (60-80 day old) and young adult mice (30-40 day old). We found that there was a significant difference in the effect of Δt +10ms STDP induction in these two groups of mice as seen in the EPSPs rising slope (Fig 1 A and B, mature adult: 53.6 ± 8.4%, n = 8; adult: 107.7 ± 2.9% vs 100 %, n = 4) and peak amplitude (Figure Supplement 1). It is important to notice that our result of lack of lasting changes of EPSPs in adult mice (30-40 day old) reproduces the previously reported lack of lasting changes of EPSPs observed in 30-50 day old mice in similar experimental conditions, i.e. EPSPs evoked in layer 5 pyramidal neurons with intact synaptic inhibition ^15^.

### Dopamine application blocks t-LTD in mature adult mice

After determining that STDP induction has an age dependent effect on EPSPs, we sought to investigate whether bath application of a cortical neuromodulator such as DA would modulate the STDP effect towards t-LTP in mature adult mice (60-80 day old) as previously shown in younger mice ^15,19^. DA (20 μM) was applied for ten minutes starting during the last minute of baseline recordings and lasting throughout the induction protocol. We found that the EPSPs recorded from cells where DA was applied during the Δt +10 ms STDP induction protocol had a significant steeper rising slope (Fig 2 A and B) and amplitude (Figure Supplement 3, panels A and B) than those recorded from cells that underwent the induction protocol in the absence of DA. As a result, DA changed the t-LTD seen in mature adult mice to no change in EPSP rising slope (Fig 2 A and B, 99.8 ± 6.7, *p* = 0.9 vs 100 %, n = 8) and peak amplitude (Figure Supplement 3, panels A and B).

**Fig 2.**
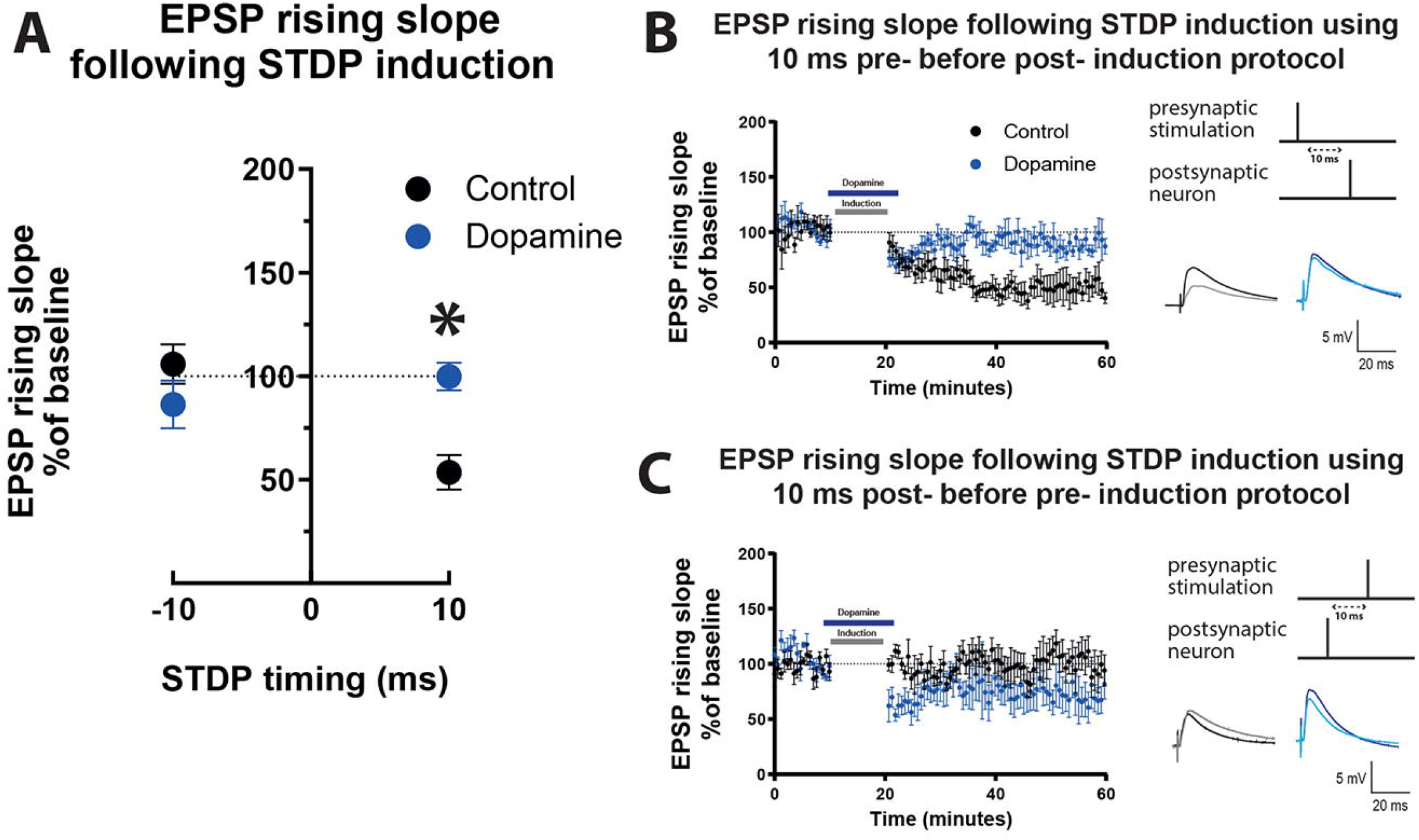
Effect of dopamine on EPSP rising slope following STDP induction in mature adult mouse cortical layer 5 pyramidal neurons. (A) STDP induction at Δt −10 ms and + 10 ms EPSP-AP pairing timings with and without 20 μM dopamine application. EPSP rising slope with the −10ms timing was not different between the control and dopamine group (*p* = 0.2). At the +10 ms timing, EPSP rising slope showed a significant change (*p* = 0.0007) from t-LTD to no change. The time-course of the EPSP rising slope during the STDP experiment using the +10 ms timing is shown in (B) and the −10 ms timing in (C). Dopamine bath application and the time of STDP induction are indicated by bars in the graphs. To the right, the STDP induction protocol timing is illustrated and below are example traces of EPSPs. The baseline trace is the darker trace, the resultant trace following STDP induction is the lighter trace; each trace is the average of 80 traces from the same recording. All data are shown as mean ± SEM. For the −10ms timing n = 6 and for the +10 timing n=8.

In contrast, when DA was applied during the Δt −10 ms STDP induction protocol, EPSP rising slope (Fig 2 A and C, 86.4 ± 11.5 vs 100 %, *p* = 0.3, n = 6) and peak amplitude (Figure Supplement 3, panels A and C) remained unchanged and there was no difference in EPSP rising slope (Fig 2 A and C), or peak amplitude (Figure Supplement 3, panels A and C) before and after DA. Furthermore, DA (20 μM) application did not affect the rising slope of EPSPs evoked at 0.14 Hz with no STDP induction protocol (Figure Supplement 4), and did not significantly affect AP firing frequency and basic electrophysiological properties (Figure Supplement 4).

The results show that DA modulates STDP by blocking t-LTD without broadening its timing window in mature adult mice.

### Optogenetically triggered release of dopamine also blocks t-LTD

Bath application of drugs can be not entirely representative of physiological conditions. In order to test the effect by DA using more physiologically relevant conditions, we expressed channel rhodopsin (ChR2) in the ventral tegmental area (VTA) dopaminergic neurons using Dat^IREScre^ mice (Fig 3A). Expression of ChR2 was confirmed by patching fluorescently identified VTA cells and stimulating with blue light pulses as in Figure Supplement 5, panel A and B. After six weeks of transfection, fibers from VTA neurons were visible in the prefrontal cortex (Figure Supplement 5, panel C). We determined that EPSP rising slope in layer V pyramidal neurons in the prefrontal cortex was not affected by blue light stimulation alone (Figure Supplement 5, panel D). To confirm DA release from fibers we analyzed afterhyperpolarization (AHP) area occurring after 5 APs evoked by a depolarizing current pulse before and after blue light stimulation. We found that the AHP area was decreased following optogenetically triggered DA release (Figure Supplement 5, panels E and F), as previously reported ^25^.

**Fig 3.**
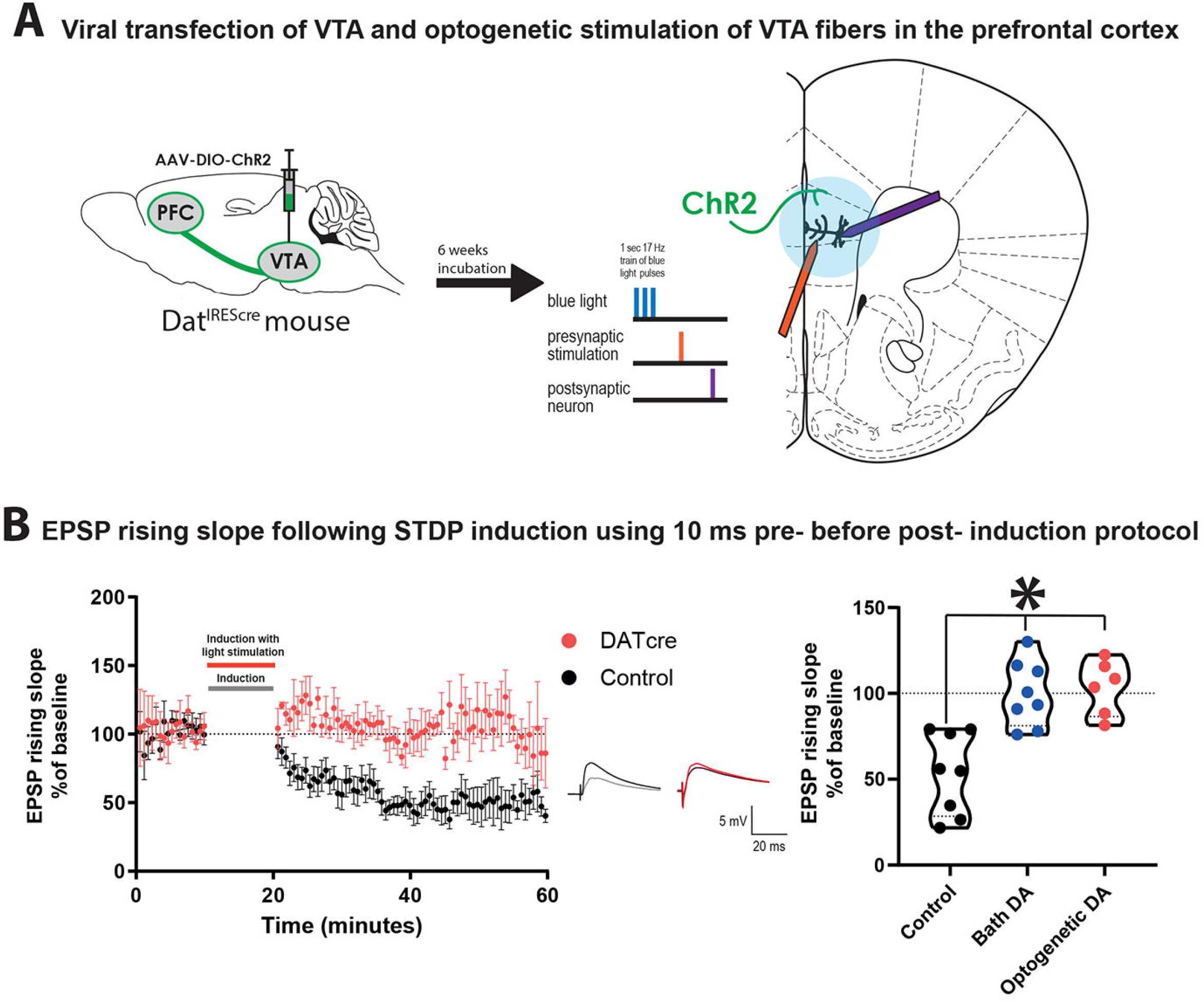
Optogenetically triggered DA release during STDP induction blocks EPSP rising time t-LTD in cortical layer 5 pyramidal neurons of mature adult mouse. (A) scheme of the viral transfection (left) and the electrophysiological protocol for STDP induction (right). Briefly, ChR2 expressing fibers from dopaminergic neuron in the VTA (shown in green) were stimulated using blue light during the STDP induction protocol. (B) The time-course of the EPSP rising slope during the STDP experiment in both control and Dat^IREScre^ mice (left) and violin plots of the summary of the data, (right). Data demonstrate that optogenetically triggered release of DA had a similar effect to bath applied DA (Tukey’s multiple comparison test, *p* = 0.99) where there is a significant difference between the control group and the DA exposed groups (Tukey’s multiple comparison test, control vs bath application of DA: *p* = 0.0006, control vs optogenetic DA: *p* = 0.0006,). In the middle example traces of EPSPs are shown. The baseline trace is the darker trace, the resultant trace following STDP induction is the lighter trace; each trace is the average of 80 traces from the same recording. All data are shown as mean ± SEM. For the optogenetic stimulation n = 6, while for the control and bath DA groups n = 8.

We then performed STDP experiments using the Δt +10 ms induction protocol on layer V pyramidal neurons in the prefrontal cortex using slices from the adult Dat^IREScre^ mice. Instead of bath application of DA, five blue light pulses were added 10 ms before the EPSP-AP pairing during the induction protocol (Fig 3A). We found that similar to bath application of DA, optogenetic DA stimulation during STDP induction also prevented the induction of t-LTD of EPSP rising slope (Fig 3B, 103.3 ± 6.5, vs 100 %, n = 6, *p* = 0.6) and peak amplitude (Figure Supplement 6). These results were significantly different from the t-LTD of the EPSP rising slope (Fig 3B, one way ANOVA, *p* = 0.0002, F = 14.3_(2,19)_) and peak amplitude (Figure Supplement 6) observed in the control group. This result suggests that bath application of DA to prefrontal cortical slices and release of DA from VTA fibers within prefrontal cortical slices evoke similar modulation on STDP of excitatory synapses in the mature adult mouse.

### Dopamine elicited t-LTP of excitatory synapses in human cortical pyramidal neurons

Next we recorded from neurons of cortical tissue resected from mature adult patients that underwent neurosurgery to test whether observations obtained in mature adult mice could be extended to humans. The patients included in the present study were on average 63.5 ±2.55 year old (Table 1). We found that the Δt +10 ms STDP induction protocol elicited a trend toward t-LTD for rising slope (Fig 4, control data, 78.5 ± 8.9, vs 100 %, *p* = 0.07, n = 5) and peak amplitude (Figure Supplement 7, control data) of EPSPs recorded from layer 5 pyramidal neurons. However, Δt +10 ms STDP induction protocol did not evoke a statistically significant change in the EPSP, in contrast to what we observed in mature adult mouse synapses (Fig 2B). Furthermore, DA bath application (20 μM) during Δt +10 ms STDP induction protocol, caused t-LTP as measured by a significant increase of the EPSP rising slope (Fig 4, DA data, 121.0 ± 6.1, vs 100 %, n = 5, *p* = 0.03) and peak amplitude (Figure Supplement 7, DA data).

**Table 1.** Basic electrophysiological properties of recorded neurons. All data is displayed as mean +/- SEM.

|  | Human | C57B/6 mice | Dat<sup>IRESc<sup>re</sup></sup> mice |
| --- | --- | --- | --- |
| **Number** | 14 | 47 | 9 |
| **Age (mouse in days,**<br>**human in years)** | 63.5 ± 2.55 | 82.5 ± 1.95 | 72.78 ± 2.77 |
| **RMP (mV)** | -68.43 ± 0.85 | -67.78 ± 0.72 | -68.91 ± 1.62 |
| **Input Resistance (MΩ)** | 119.0 ± 14.24 | 119.5 ± 5.31 | 146.0 ± 14.59 |
| **Capacitance (pF)** | 221.6 ± 21.22 | 180.7 ± 9.62 | 159.0 ± 21.49 |
| **Sag Ratio** | 1.21 ± 0.02 | 1.10 ± 0.01 | 1.09 ± 0.03 |
| **Spike Amplitude (mV)** | 105.0 ± 1.90 | 107.4 ± 1.21 | 108.5 ± 2.01 |

**Fig 4.**
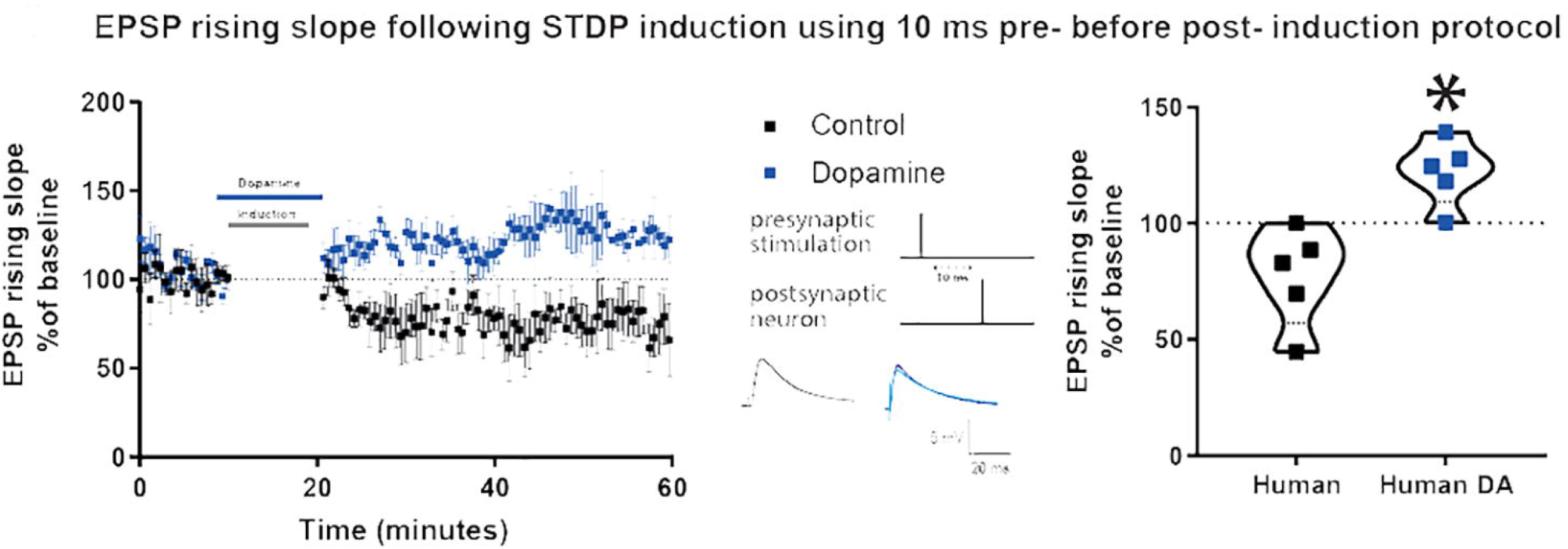
DA potentiates baseline EPSP rising time after Δt +10ms STDP protocol in adult human cortical layer 5 pyramidal neurons. Left, the time-course of the EPSP rising slope after Δt +10ms STDP induction protocol. Middle, the STDP induction protocol timing is illustrated and below are example traces of EPSPs. The baseline trace is the darker trace, the resultant trace following STDP induction is the lighter trace; each trace is the average of 80 traces from the same recording. Right, violin plots of summary of the results showing a significant difference between EPSP rising slope with and without DA application during STDP induction (*p* = 0.004). All data are shown as mean ± SEM. For both groups, n = 5.

Control experiments indicated that in the absence of STDP induction, DA application (20 μM) did not affect the rising slope of EPSPs evoked at 0.14 Hz, AP firing frequency and basic electrophysiological properties (Figure Supplement 8).

### Synaptic inhibition contributes toward different dopamine modulation of STDP at excitatory synapses of mature adult mice *versus* humans

DA evoked t-LTP at mature adult human but not at mature adult mouse excitatory synapses. What factors could contribute to this difference? DA receptors on mouse interneurons have been implicated in the mechanism of DA potentiation of STDP in younger mice ^15^. We first sought to determine if this was also true for excitatory synapses at mature adult mouse neurons. Before performing this test, we assessed whether synaptic inhibition was recruited by the stimulation protocol used and could impact EPSP kinetics. Application of the GABA-A receptor antagonist 1 μM gabazine increased EPSP peak amplitude and area, and did not change EPSP rising slope (Figure Supplement 9, panel A) recorded in mature adult mouse neurons. Then, inhibitory postsynaptic currents (IPSCs) were recorded at 0 mV, at equilibrium potential for EPSC, from layer V pyramidal neurons of mature adult mice. We found that the Δt +10 ms STDP induction protocol, delivered in current clamp at resting membrane potential, caused an increase in the IPSC rising slope (Fig 5A, 112.2 ± 3.8, vs 100 %, *p* = 0.02, n = 8) but not its peak amplitude (Figure Supplement 10A). Furthermore, DA applied during the Δt +10 ms STDP induction protocol resulted in a t-LTD for the IPSC rising slope (Fig. 5A; 87.2 ± 5.2, vs 100 %, n = 8, p = 0.04) and peak amplitude (Figure Supplement 10A). When comparing the rising slope (Fig 5A) and the peak amplitude (Figure Supplement 10A) of the IPSCs observed after STDP induction with or without DA, they were significantly different.

**Fig 5.**
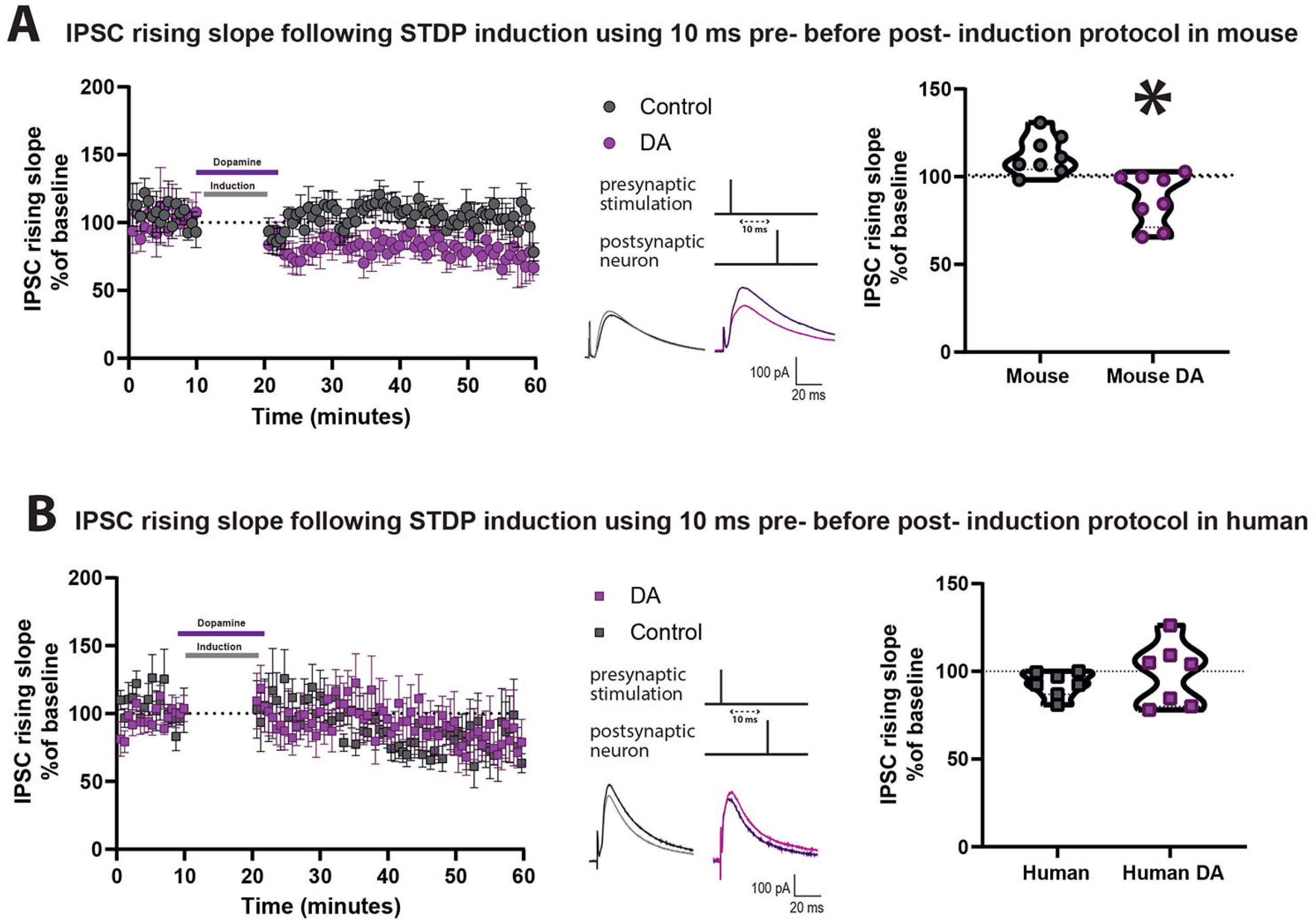
DA depresses IPSC rising slope after Δt +10ms STDP protocol in cortical layer 5 pyramidal neurons of mature adult mice but not humans. Effect of DA on IPSC rising slope following STDP induction in adult mouse (A) and human (B) pyramidal neurons. Left, the time-course of the IPSC rising slope after Δt +10ms STDP timing protocol. Middle, the STDP induction protocol timing is illustrated and below are example traces of IPSCs. The baseline trace is the darker trace, the resultant trace following STDP induction is the lighter trace; each trace is the average of 80 traces from the same recording. Right, violin plots showing summary of the results. DA significantly reduced the IPSC rising slope after STDP induction in neurons recorded from mature adult mice (*p* = 0.002, n = 8), but not from humans (*p* = 0.5, n = 7). All data are shown as mean ± SEM.

Data from the Allen Institute for Brain Science RNA-seq transcriptional profiles shows that DA receptors 1 and 2 have lower expression in human GABAergic neurons of cortical layer 5 than in mouse (Figure Supplement 11). Therefore, we also examined the effect of DA application on IPSC caused by Δt +10 ms STDP induction delivered in current clamp at resting membrane potential in human pyramidal neurons. We found t-LTD after the STDP induction protocol, as assessed by the IPSC rising slope (Fig 5B, 92.9 ± 2.7, vs 100 %, n = 7, *p* = 0.04) and a non-significant trend towards t-LTD in peak amplitude (Figure Supplement 10B). Furthermore, we observed that there was no change in IPSC rising slope (Figure 5B, 98.2 ± 6.7, vs 100 %, n = 7, *p* = 0.9) and peak amplitude (Figure Supplement 10B) in neurons exposed to the Δt +10 ms STDP induction protocol in the presence of 20 μM DA. When comparing the rising slope (Fig 5B) and the peak amplitude (Figure Supplement 10B) of the IPSCs observed after STDP induction with or without DA, they were not significantly different.

Control experiments indicated that DA alone had no effect on the rising slope of IPSCs evoked by 0.14 Hz stimulation in the absence of the STDP induction protocol in both human and mouse neurons (Figure Supplement 12). We also confirmed that the IPSCs recorded were entirely mediated by GABA-A receptors, as application of 1 μM gabazine abolished the evoked IPSCs, as shown in examples in human and mouse neuron recordings in Figure Supplement 9, panel B.

Our results suggest subtle differences between STDP of synaptic inhibition and its DA modulation at layer 5 pyramidal neurons in mature adult mice compared to mature adult human patients. These differences could represent one of the mechanisms underlying the different effects by STDP of excitatory transmission and its modulation by DA in mature adult mouse *versus* adult human patients.

## Discussion

The key findings of the present study can be summarized as following. We observed an anti-Hebbian t-LTD triggered by one post-synaptic AP preceded by pre-synaptic spiking by 10ms (Δt +10ms) evoked by extracellular stimulation with intact synaptic inhibition in layer 5 pyramidal neurons of the neocortex from mature adult mice *in vitro*. Exogenous application of DA or optogenetic stimulation of VTA fibers, to release endogenous DA, switched this t-LTD into no change in the EPSP after the pairing protocol. When a burst of APs (and not only one AP) were evoked in the post-synaptic neuron during the pairing protocol, then a stable EPSP (i.e. no t-LTD) was observed, presumably by transiently boosting post-synaptic dendritic calcium via back-propagating APs. Furthermore, we also investigated STDP and DA modulation in cortical pyramidal neurons from mature adult neurological patients using the same experimental conditions used in mice. We observed that the Δt +10ms protocol elicited a trend toward t-LTD, although this synaptic depression was not statistically significantly different from baseline. Application of DA strengthened the EPSP toward t-LTP. We also observed that in mature adult mice, a STDP protocol enhanced synaptic inhibition but this effect was reversed to t-LTD after DA application. In contrast, in adult humans, a STDP protocol without or with DA evoked no change of synaptic inhibition.

Using classical STDP pairing protocols, we observed t-LTD after the Δt +10ms when synaptic inhibition was left unaffected in mature adult mice (60-80 day old). In contrast, the same pairing protocol performed on juvenile excitatory synapses usually evoke t-LTP in the neocortex of rodents ^21,26^. Importantly, when we tested younger mice (range 30-50 day old same as in ^15^), we observed no lasting changes in the EPSP following the Δt +10ms protocol, exactly as previously reported in similar experimental conditions ^15^. The detection of different plastic rules between young adult mice (30-50 day old) and mature adult mice (60-80 day old) can be surprising, because a month old mouse is often assumed to reach adulthood and show constant features ^27^. However, young and mature old mice can also express important differences. For example, 30 day old and 70 day old mice with deletion of the NMDA receptor NR1 subunit gene in glutamatergic neurons display different social behavior ^28^. Furthermore, 30 day *versus* 70 day old neuroligin-3 knock in mice show significant differences in the volume of cortical white and gray matter associated with altered sociability ^29^.

Our data suggest that when STDP is tested under intact GABAergic inhibition, t-LTD is the prominent form of synaptic plasticity that occurs in mature adult rodent neocortex, in contrast to what have been seen at juvenile synapses showing t-LTP ^21,26^. As cortical layer V neurons of adult, neurons from mature mice are under powerful constraint by local GABAergic interneurons, which may explain the lack of t-LTP induced by a mild induction STDP protocol. When we applied a burst of postsynaptic APs and not only one AP in the STDP induction protocol a stable EPSP without t-LTD was observed. Our result is consistent with the assumption that in older animals, synapses tend to require stronger induction protocols for t-LTP to occur ^24^. Of note, Meredith et al^24^ studied neocortical synapses of rodents (up to 45 days old), an age range comparable to the study published by ^15,16^, but younger than our mature adult mouse sample.

Our results of t-LTD in mature adult mice differs from some previous reports performed in the rat, that suggested that the capacity of rodent cortical synapses to undergo STDP t-LTD decline with age ^23,30^. The difference could be explained by many factors including different species: rat *versus* mouse; age: mature adult *versus* adult or juvenile animals; cortical areas tested: barrel cortex *versus* parietal, prefrontal and temporal cortex; neuron types: recordings in neurons with the soma in layer 2-4 ^30^ *versus* layer 5 (present study).

Our results obtained in mature synapses triggered by pre before-post spike pairings formally resemble anti-Hebbian STDP observed in some juvenile excitatory cortical synapses. For example, pre < 25 ms before-post spike pairings of spiny stellate neurons in layer 4 of the barrel field in young rat somatosensory cortex elicits t-LTD ^13^. A general explanation for the presence of both Hebbian and non-Hebbian t-LTD at various cortical excitatory synapses have been given some time ago ^31–33^. According to this view, the sign of synaptic plasticity in neocortical pyramidal neurons is regulated by the spread of APs to the synapse. This creates a progressive gradient between t-LTP and t-LTD as the distance of the synaptic contacts from the soma increases. Furthermore, we observed a lack of STDP either after pre-before-post spike pairings as well as post-before-pre pairings with a longer time interval of +/- 30 ms in mature adults mice 60-80 days old consistent with previous results obtained in young adult mice (30-50 days old) ^16^.

Regarding our data obtained at excitatory synapses of neocortical human cortex, they are consistent with a study documenting STDP on pyramidal neurons of layers 2-6 performed in acute slices of human cortex resected from tumor or epilepsy patients ^23^. As in our sample, this study reported that multiple pre-before-post spike pairings (especially short intervals 5-10 ms) evoke STDP t-LTD, confirming anti-Hebbian STDP rules at cortical excitatory synapses. In contrast to our data however, this study also found that post-before-pre pairings (especially at short intervals 5-10 ms) evoke STDP t-LTP. Different experimental conditions (e.g., age and patient conditions and cortical areas studied) could account for these discrepancies.

Our results show that DA promoted synaptic strengthening in both mouse and human cortical excitatory synapses, but t-LTP was observed only at human cortical synapses and not at mouse cortical synapses where synaptic responses remained close to the baseline after the Δt +10ms protocol. Our observations are well aligned with the idea that DA controls the polarity of STDP ^21^. For example, DA applied during STDP induction leads to t-LTP with spike timing that would induce t-LTD in control conditions ^14,19^. This modulatory effect by DA was also observed in layer 5 pyramidal cells of the PFC tested in 30-50 days old mice ^16^. Previous work also reported some differences between data obtained in rodents and in human cortex. For example, as already discussed above, Verhoog et al.^23^ observed only t-LTP in rodents for both negative and positive pre- and post-synaptic timing intervals, whereas pre before-post spike pairings evoke t-LTD and post-before-pre pairings elicit t-LTP at human cortical synapses. Moreover HCN1-channel-related gene expression and function is more prominent in human than mouse supragranular cortex ^34^. This difference generates peculiar synaptic integration in human supragranular pyramidal neurons that could affect the effects of STDP.

Many presynaptic and/or postsynaptic factors could account for species-specific differences we observed. We have identified one of them, namely a differential impact onto inhibitory interneurons presynaptic to layer 5 pyramidal cells. Our data show that STDP induction alone potentiated IPSCs recorded from mature adult mice. We also observed that the peak amplitude of EPSPs overlapped with the initial part of the IPSP and it was sensitive to a GABA-A receptor antagonist. Therefore, this potentiation of synaptic inhibition could contribute to the t-LTD effect found in the EPSPs recorded in mature adult mice. Furthermore, application of DA during a STDP protocol depressed inhibitory transmission in the cortex of mature adult mice. This suggests that DA applied during a STDP protocol may exert at least some of its effect by disinhibition resulting in t-LTD of EPSP being converted to no change of synaptic strength in mature adult mice. This interpretation is consistent with the idea that DA type 2 receptors expressed by cortical inhibitory axonal terminals mediate the allosteric inhibition of GABA release and contribute to DA STDP gating in young adult mice ^15^. In the human pyramidal neurons, STDP induction alone had no effect on IPSCs and we did not see a t-LTD effect in the EPSP such as the one seen in the mature adult mouse. Furthermore, when DA was applied during a STDP protocol, there was again no change in the IPSC, however we observed a potentiation in the EPSP, suggesting that presynaptic inhibition is not a target for DA modulation at human cortical circuits. Our data are consistent with data from the Allen Institute for Brain Science showing that DA receptors 1 and 2 have lower expression in human layer 5 cortical GABAergic neurons than in mouse.

Several other mechanisms may account for the species-specific differences we have observed. Amongst them, a key postsynaptic factor could be represented by the stronger compartmentalization of cortical layer 5 pyramidal neurons in humans than rodents ^35^, a feature that could be under neuromodulatory control. Ongoing and future work will provide more details on the specializations of cortical pyramidal neurons in humans as compared to rodents ^36–38^. They could lead to identification of differences in the integration of back-propagating APs and synaptic inputs resulting in human neurons specific STDP phenotypes.

Finally, it is important to acknowledge some methodological limitations present in our study. Firstly, the extracellular stimulation method used precluded any defined information on the microcircuit investigated. During the experiments, the stimulation electrode was placed at layer 2-3, about 100-150 μm more superficial than the recording electrode placed in layer 5. Therefore, the stimulation electrode activated a heterogeneous set of extrinsic excitatory fibers (e.g., interhemispheric corticocortical afferents) and intrinsic axons (e.g., collaterals of pyramidal neurons). Moreover, the interneuron types directly or indirectly activated by the stimulation remained undefined ^39^. Secondly, it is important to admit that the use of tissue from human cerebral cortex of patients subjected to neurosurgery has some intrinsic methodological limitations. One of these is that the tissue may have some pathological features that remain undetected. We have used cortical tissue from patients with low grade glioma tumour and although we recorded only from neurons located outside the main tumor, some infiltration cannot be excluded. Another possible limitation is the variability due to heterogeneity of cortical areas of provenance, different age and sex of the patients, their individual clinical and pharmacological history (see Table 2). Despite this variability, we observed basal functional parameters that were rather homogeneous across samples and patients and homogeneity in the effects mediated by DA, consistent with other reports testing pharmacological agents in human cortical slices ^40–42^. Thirdly, we recorded from layer 5 pyramidal neurons and pooled the data as these neurons were an homogeneous group. Recent data clearly indicate a pyramidal neuron type diversity based on genetic profile expression and projection sites ^36^, but we did not attempt to take this aspect into account, as this would have been beyond the scope of our study. Fourthly, DA gating has been associated with reward and prediction errors as well as novelty and salience detection ^43^. We assume that the STDP model represents a cellular correlate underlying cognition, but how good is this assumption? One of the key unsolved issues is represented by the so-called distal reward problem, that is how neuronal networks, despite a temporal gap, identify which past networks activities led to reward and which are irrelevant ^44^. One way to help solving this problem would be the identification of an eligibility trace generated by the STDP spiking activity and triggered by DA ^45^, but this hypothesis awaits experimental demonstration.

**Table 2.** Donor patients attributes.

|  | **Age** | **Sex** | **Histological diagnosis, tumor grade** | **Tumor IDH** | **Tumor localization, lateralization** | **Epilepsy, type** | **Anaesthesia** |
| --- | --- | --- | --- | --- | --- | --- | --- |
| 1 | 56-60 | Male | Glioblastoma, 4 | Wild Type | Parietal, right | Yes, focal | Prednisolone, Dexamethasone, Metaxedrine /Phenylephrine, Remifentanyl, Propofol |
| 2 | 76-80 | Male | Glioblastoma, 4 | Wild Type | Temporal, right | No | Dexamethasone, Metaxedrine /Phenylephrine, Ephedrine, Remifentanyl, Fentanyl, Propofol |
| 3 | 70-75 | Female | Tissue necrosis | N/A | Temporal, right | No | Dexamethasone, Metaxedrine/ Phenylephrine, Remifentanyl, Propofol |
| 4 | 50-55 | Male | Glioblastoma, 4 | Wild Type | Temporal, right | Yes focal | Dexamethasone, Metaxedrine/ Phenylephrine, Remifentanyl, Propofol |
| 5 | 60-65 | Male | Glioblastoma, 4 | Wild Type | Temporal, left | No | Prednisolone, Noradrenalin, Metaxedrine /Phenylephrine, Ephedrine, Remifentanyl, Propofol |
| 6 | 60-65 | Female | Glioblastoma, 4 | Wild Type | Occipital, right | Yes, generalized | Dexamethasone, Noradrenalin, Metaxedrine /Phenylephrine, Ephedrine, Remifentanyl, Fentanyl, Propofol |
| 7 | 70-75 | Male | Glioblastoma, 4 | Wild Type | Temporal, right | No | Prednisolone, Metaxedrine/ Phenylephrine, Ephedrine, Remifentanyl, Propofol |
| 8 | 70-75 | Female | Glioblastoma, 4 | Wild Type | Frontal, left | No | Dexamethasone, Metaxedrine/Phenylephrine, Ephedrine, Remifentanyl, Fentanyl, Propofol |
| 9 | 60-65 | Female | Glioblastoma, 4 | Wild Type | Parietal, left | No | Dexamethasone, Metaxedrine/Phenylephrine, Remifentanyl, Propofol |
| 10 | 40-45 | Male | Glioblastoma, 4 | Wild Type | Parietal, left | No | Dexamethasone, Metaxedrine/Phenylephrine, Remifentanyl, Fentanyl, Alfentanyl, Propofol |
| 11 | 56-60 | Male | Glioblastoma, 4 | Wild Type | Temporal, left | Yes, generalized | Dexamethasone, Metaxedrine/Phenylephrine, Ephedrine, Remifentanyl, Fentanyl, Alfentanyl, Propofol |
| 12 | 56-60 | Male | Glioblastoma, 4 | Wild Type | Temporal, left | Yes focal | Metaxedrine/Phenylephrine, Ephedrine, Remifentanyl, Fentanyl, Propofol |
| 13 | 60-65 | Male | Glioblastoma, 4 | Wild Type | Temporal, right | No | Prednisolone, Dexamethasone, Metaxedrine/Phenylephrine, Ephedrine, Remifentanyl, Propofol |
| 14 | 70-75 | Female | Glioblastoma, 4 | Wild Type | Temporal, right | No | Dexamethasone, Metaxedrine/Phenylephrine, Ephedrine, Remifentanyl, Fentanyl, Propofol |

Despite these limitations, our findings, and particularly the discovery of DA modulation of STDP at excitatory synapses of layer 5 neurons in human cortex, provide information that can be further explored by future experiments. For example, the cellular correlates of DA-mediated gating of cognitive process in the human cortexcould be explored by using an in *situ* multi-electrode array recording approach.

## Methods

### Animals

C57B/6J mice and DAT^IREScre^ mice were purchased from Jackson Laboratory and bred in-house in the animal facility of the Department of Biomedicine, Aarhus University. Mice were group housed in a temperature and humidity controlled plastic vivarium in a 12 hour regular light/dark cycle with lights on at 8:00 a.m. All procedures with animals were approved by and conducted in accordance to with the The Animal Experiments Inspectorate under the Ministry of Environment and Food of Denmark (License number 2017-15-0201-01201).

### Viral transfection

Nine DAT^IREScre^ mice (P35-45) form four litters were anaesthetized using a mix of 0.05 mg/ml of Fentanyl (Hameln pharma ltd, UK;0.05 mg/kg), plus 5 mg/ml of Midazolam (Hameln Pharma Ltd, UK;) 5 mg/kg) and 1 mg/ml of medetomidine (0.5 mg/kg; VM Pharma). They then received bilateral injections of ssAAV-9/2-hEF1α-dlox-hChR2(H134R)_EYFP(rev)-dlox-WPRE-hGHp(A) (1-2 μL; titre: 6.0 x 10^12 vg/ml; VVF, Switzerland) into the VTA at the following coordinates: anterior-posterior +3.1mm, medio-lateral +/- 0.5mm, dorsal-ventral −4.5mm with respect to Bregma. Injections were made through a pulled 1mm glass pipette using a Picospritzer III (Parker Hannifin, US). Pipettes were kept in place for at least 10 minutes after injection of the virus. Following surgery, animals received 0.1mg/kg of Buprenorphine (Temgesic; Indivior UK Limited) subcutaneously and an antidote mix of 0.4 mg/ml Naloxone (B. Braun, 115241) 1.2 mg/kg), plus 5 mg/ml of Atipamelozone Hydrochloride (2.5 mg/kg) and 0.5mg/ml of Flumazenil ((Hameln Pharma Ltd, UK) 0.5 mg/kg) to reverse the anesthesia. Then, mice were single-housed. We recorded from VTA neurons three weeks after surgery from two animals. Furthermore, we recorded neurons in the prefrontal cortex six weeks after surgery from seven animals. This extra time was necessary to be able to detect the virus expression in fibres of the prefrontal cortex (Figure Supplement 5, Panel C).

### Human tissue acquisition

All procedures with human tissue and data were approved by and conducted in accordance with the The Scientific Ethics Committee for the Region of Midtjylland Denmark (official name in Danish: De Videnskabsetiske Komitéer for Region Midtjylland) (project number M-2017-82-17). All patients provided informed consent for tissue donation. Human brain tissue samples were obtained from Aarhus University Hospital in collaboration with their neurosurgery team. Patients in this study were undergoing surgery for a deep brain tumour and the samples we received were from surgically excised tissue that needed to be removed in order to gain access to the tumour. The sample provided was taken as far from the tumour as was feasible, this was typically 5-10 mm. Samples were mainly taken from the temporal lobe and in some cases the parietal, occipital and frontal lobes. See table 2 for more detailed information of the patients involved in this study.

Human brain tissue samples were surgically excised and immediately placed in ice-cold *N*-methyl-D-glucamine (NMDG)-based artificial cerebral spinal fluid (aCSF). NMDG aCSF was prepared as previously described ^46^, the composition in mM was 92 NMDG, 2.5 KCl, 1.25 NaH2PO4, 30 NaHCO3, 20 4-(2-hydroxyethyl)- 1-piperazineethanesulfonic acid (HEPES), 25 glucose, 2 thiourea, 5 Naascorbate, 3 Na-pyruvate, 0.5 CaCl2·4H2O and 10 MgSO4·7H2O. The pH was titrated to 7.3–7.4 with hydrochloric acid and the osmolality was 300–310 mOsmoles/Kg. The solution was chilled on ice and bubbled with carbogen gas. Once the tissue was taken out of the operating room, we removed any excess blood and white matter before placing the sample into a fresh tube of oxygenated NMDG aCSF. The sample was placed on ice, connected to a portable container of carbogen gas and transported to the laboratory (approximately 15 minutes travel time).

### Acute ex vivo brain slice preparation

Mice were killed by decapitation while under isoflourane anaesthesia. The brain was removed and placed in ice-cold NMDG based aCSF for approximately two minutes. Mouse brains were blocked and mounted on the vibrating microtome platform to cut coronal sections of the temporal cortex, parietal cortex, prefrontal cortex or VTA. Human brain samples were mounted such that the blade was perpendicular to the pial surface. We did not remove the pia mater. From this point, the procedure for preparation of adult mouse and human acute ex vivo brain slices was the same. Acute brain slices 350 μm in thickness were sliced on a Leica 1200S vibrating mictrome (Leica Microsystems, Denmark). Slices were then placed in a recovery chamber containing 32°C, carbogenated NMDG aCSF for 12 minutes. Slices were then transferred to a holding chamber with room temperature carbogenated aCSF composed (in mM) of: 92 mM NaCl, 2.5 mM KCl, 1.25 mM NaH2PO4, 30 mM NaHCO3, 20 mM HEPES, 25 mM glucose, 2 mM thiourea, 5 mM Na-ascorbate, 3 mM Na-pyruvate, 2 mM CaCl2·4H2O and 2 mM MgSO4·7H2O, with a pH of 7.3, and osmolarity of 300-310 mOsmoles/Kg. Slices were left to recover for at least one hour before recording and were stored for up to 12 hours.

### Electrophysiology

Brain slices were transferred to a recording chamber mounted on a SliceScope microscope (Scientfica, UK) and superfused with carbogenated ACSF composed of 119 mM NaCl, 2.5 mM KCl, 1.25 mM NaH2PO4, 24 mM NaHCO3, 12.5 mM glucose, 2 mM CaCl2·4H2O and 2 mM MgSO4·7H2O. Pyramidal cells in layer V of the mouse or human temporal, parietal or prefrontal cortex were visualized with infrared differential interference contrast microscopy. A SciCam Pro camera (Scientifica, UK) was used for visualization and image capture. Whole-cell recordings were performed using borosilicate glass pipettes pulled with a horizontal pipette puller (DMZ universal electrode puller, Zeitz, Germany). Pipettes contained intracellular solution consisting of 126 mM K-gluconate, 10 mM HEPES, 4mM KCl, 4 mM Mg-ATP, 0.3 mM Na3-GTP and 10 mM Na2-phosphocreatine. Liquid junction potential was calculated to be −16 mV and was not corrected for. For recordings of IPSCs only, intracellular solution consisted of 65mM CsMeSO3, 65 mM K-gluconate, 10mM HEPES, 10 mM CsCl2, 4mM MgCl, 0.1 mM EGTA, 2 mM Mg-ATP, 0.3 mM Na3-GTP, 10 mM Na2-phosphocreatine. Liquid junction potential was calculated to be −11 mV and was not corrected. For both solutions the pH was 7.3-7.4 and 290 mOsmoles/Kg. When the electrodes were filled with an internal solution had an estimated resistance ranging 3-5 MΩ. Recordings were made using a Multiclamp 700B amplifier, acquired at 20 kHz, low-pass filtered at 2 kHz using a Digidata 1550B low noise data acquisition system (Molecular Devices, USA). For whole-cell recordings, pipette capacitance was neutralized and bridge balance applied. Recorded cells had an initial resting membrane potential between −60 and −75 mV. Recordings were included only if they had a change in series resistance of <25%. A baseline recording was obtained for at least ten minutes. All experiments were performed in the absence of GABAergic transmission blockers.

### Spike timing dependent plasticity protocol

A stimulating electrode inserted in a glass pipette filled with recording aCSF with a resistance of approximately 1 MΩ was placed approximately 150 μm from the soma nearby the apical dendrite. As described in Verhoog et. al.^23^, EPSPs with an amplitude between 3-8 mV or IPSCs with an amplitude between 100-300 pA were evoked at a rate of 0.14 Hz (stimulation parameters were 100 μs duration and 200-500 μA intensity) controlled by an A360 stimulus isolator (World Prescision Instruments, UK). Cells were held between −68 to −72 mV for evoked EPSPs and 0 mV for IPSCs.

Pairing was always conducted in current clamp mode, for both EPSPs and IPSPs. Various pairing protocol timings were used, from 30 ms pre- before post-synaptic stimulation to 30 ms post-before pre- synaptic stimulation. For all timings, an induction protocol of 75 pairings of the EPSP with an AP generated by direct stimulation to the cell body (5 ms duration, intensity between 500 - 1500 pA) a rate of 0.14 Hz was used. For the burst pairing protocol, a 25 ms depolarization with an intensity between 500 – 100 pA was used to induce 3-4 APs. Dopamine (20 μM, Tocris) was bath applied for ten minutes starting during the last minute of baseline recordings and lasting throughout the induction protocol.

Following the induction protocol, EPSP/IPSCs were evoked at a rate of 0.14 Hz, as they were during the baseline recording. They were then monitored for the next 40-50 minutes for analysis. The EPSP rising slope (20-80%) and peak amplitude used for analysis were calculated from a five minute most stable section (defined as section that best matched baseline RMP and series resistance levels) between 25 - 35 minutes after the induction protocol.

### Pharmacology

To confirm that evoked IPSCs were, in fact, GABAergic, 1 μM gabazine (SR 95531, Tocris) was applied at the end of an STDP recordings. To see the effect of GABAergic synaptic transmission on EPSP kinetics, 1 μM gabazine was applied for 5 minutes to a separate set of mouse neurons while evoking EPSPs at a rate of 0.14 Hz, as described above.

To see the effect of dopamine bath application on EPSP kinetics and basic cellular properties, 20 μM dopamine was prepared in a light protected beaker to prevent oxidation by light. Dopamine was then applied for 5 minutes to mouse and human neurons while evoking EPSPs at a rate of 0.14 Hz, as described above.

### Optogenetics

For DAT^IREScre^ mice, endogenous release of dopamine during the induction protocol was achieved by stimulating ChR2 in the fibers of VTA projections. These fibers were located by eYFP expression. Terminals were stimulated with blue light from a CoolLed PE-300ultra (Scientifica) (460 nm, ~10mW power). Each stimulation consisted of a 1 second, 17 Hz train of blue light pulses with a 5ms pulse width, 10 ms before the pairing of the EPSP and AP during the induction protocol.

To confirm endogenous dopamine release in the prefrontal cortex, we assessed the effect of blue light stimulation of ChR2 on the layer V pyramidal neuron AHP as described in ^25^. In brief, 5 APs were generated with a 60 ms pulse of 1000-1500 pA and AHP area was analyzed before and after seven minutes of repeated blue light exposure, 1 second of 17 Hz train at 0.14Hz to mimic dopamine release caused during the induction protocol. All these protocols of optogenetic stimulations were adopted from Buchta et al ^25^, who also investigated the action of DA released from VTA terminals in rodent PFC. We also stimulated VTA neurons, identified by eYFP expression, using a 10Hz pulse train (10 ms pulse width) of blue light (Figure Supplement 5, panels A and B) to confirm expression of ChR2.

### Quantification of dopamine receptor expression

Using publicly available data from the Allen Institute for Brain Science, we extrapolated the single nucleus mRNA expression data for dopamine receptor 1 and 2 (DRD1 and DRD2, respectively). The database used can be found at https://portal.brain-map.org/atlases-and-data/rnaseq and detailed methods of single nuclei fluorescence-activated cells sorting (FACS) isolation followed by Smart-seq v4 based library preparation and single-cell deep (2.5 million reads/cell). RNA-Seq can be found in ^36^. Data was derived from human medial temporal gyrus tissue samples and adult mouse primary visual cortex and anterior lateral motor area (postnatatal days 53-59). The data was organized by cortical layer expression in GABAergic neurons. Plots were made using GraphPad prism and represent the distribution of mRNA expression on a log scale of counts per million (CPM).

### Data analysis

All electrophysiological data was acquired using pCLAMP 10.7 and analyzed in clampfit 10.7 (Molecular Devices, US). Statistical analysis was performed using GraphPad Prism 8 (GraphPad Software, USA). All EPSP/IPSC rising slopes and amplitudes, and basic electrophysiological properties were normalized to baseline for analysis. All data are presented as mean ± SEM. Due to the nature of whole-cell electrophysiology and availability of human tissue, there is a limited sample size. We have run statistics with both parametric and nonparametric measures and found virtually no difference. The Shapiro-Wilk test for normality can be run on small data sets (as low as n = 3) ^47^ and we found that all data was normally distributed unless otherwise indicated in the text. Therefore, we present the following: when comparing two groups, Student’s *t* test was used, when comparing the effect of STDP to baseline a one sample *t* test was used, when more than two groups were compared, an ANOVA was used. Sample sizes refers to the number of recorded neurons. No more than two neurons using the same protocol were sampled from the same mouse or human brain sample.

## Acknowledgements

We thank Lise Moberg Fitting: she collected human cortical tissue samples from the neurosurgery operating room and transported it to a room nearby where the tissue was immediately placed in ice-cold NMDG aCSF; she also compiled patients’ clinical data. We also thank Majken Sand for her help on human cortical tissue transport from Aarhus University hospital to Aarhus University lab and for her help with the preparation of aCSF and drug aliquots. We acknowledge Dr. Wen-Hsien Hou and Meet Jariwala for their assistance during the human tissue sample collection. We thank Prof. Peter Somogyi, Dr. Martin Field and Istvan Lukacs, Oxford University, Dr. Wen-Hsien Hou and Dr. Sadegh Nabavi, Aarhus University, for helpful comments on this manuscript.

This work was supported by an ERC grant (ERC-2015-AdG 694988 to M.C.). We acknowledge the support of the Natural Sciences and Engineering Research Council of Canada (NSERC), [PDF-532810-2019 to E.L.L.].

## Author contributions

E.L.L., conceptualization, investigation, analysis, writing; R.L.J., neurosurgery; commenting text; A.R.K., neurosurgery, application to use human patients sample for research; J.C.H.S, neurosurgery supervision, funding acquisition, commenting text; M.C., conceptualization, writing, funding acquisition, supervision.

## Additional information

The authors declare no competing interests.

## Notes

### Competing Interest Statement

The authors have declared no competing interest.

## References

1. Arnsten, A. F. T., Wang, M. J. & Paspalas, C. D. Neuromodulation of Thought: Flexibilities and Vulnerabilities in Prefrontal Cortical Network Synapses. Neuron 76, 223–239 (2012).

2. Thiele, A. & Bellgrove, M. A. Neuromodulation of Attention. Neuron 97, 769–785 (2018).

3. Klanker, M., Feenstra, M. & Denys, D. Dopaminergic control of cognitive flexibility in humans and animals. Front. Neurosci. 7, 1–24 (2013).

4. Ott, T. & Nieder, A. Dopamine and Cognitive Control in Prefrontal Cortex. Trends Cogn. Sci. 23, 213–234 (2019).

5. Björklund, A. & Dunnett, S. B. Dopamine neuron systems in the brain: an update. Trends Neurosci. 30, 194–202 (2007).

6. Merten, K. & Nieder, A. Active encoding of decisions about stimulus absence in primate prefrontal cortex neurons. Proc. Natl. Acad. Sci. U. S. A. 109, 6289–6294 (2012).

7. Schultz, W., Dayan, P. & Montague, P. R. A neural substrate of prediction and reward. Science (80-.). 275, 1593–1599 (1997).

8. Bi, G. Q. & Poo, M. M. Synaptic modifications in cultured hippocampal neurons: dependence on spike timing, synaptic strength, and postsynaptic cell type. J. Neurosci. 18, 10464–10472 (1998).

9. Debanne, D., Gähwiler, B. H. & Thompson, S. M. Long-term synaptic plasticity between pairs of individual CA3 pyramidal cells in rat hippocampal slice cultures. J. Physiol. 507, 237–247 (1998).

10. Markram, H., Lübke, J., Frotscher, M. & Sakmann, B. Regulation of synaptic efficacy by coincidence of postsynaptic APs and EPSPs. Science 275, 213–5 (1997).

11. Miller, K. D., Abbott, L. F. & Song, S. Competitive Hebbian learning through spike-timing-dependent synaptic plasticity. Nat. Neurosci. 3, 919–926 (2000).

12. Hebb, D. O. The Organization of Behavior; A Neuropsychological Theory. (J. Wiley; Chapman & Hall, 1949). doi: 10.2307/1418888.

13. Egger, V., Feldmeyer, D. & Sakmann, B. Coincidence detection and changes of synaptic efficacy in spiny stellate neurons in rat barrel cortex. Nat. Neurosci. 2, 1098–1105 (1999).

14. Zhang, J.-C., Lau, P.-M. & Bi, G.-Q. Gain in sensitivity and loss in temporal contrast of STDP by dopaminergic modulation at hippocampal synapses. Proc. Natl. Acad. Sci. 106, 13028–13033 (2009).

15. Xu, T. X. & Yao, W. D. D1 and D2 dopamine receptors in separate circuits cooperate to drive associative long-term potentiation in the prefrontal cortex. Proc. Natl. Acad. Sci. U. S. A. 107, 16366–16371 (2010).

16. Ruan, H., Saur, T. & Yao, W. D. Dopamine-enabled anti-Hebbian timingdependent plasticity in prefrontal circuitry. Front. Neural Circuits 8, 1–12 (2014).

17. Edelmann, E. & Lessmann, V. Dopamine modulates spike timingdependent plasticity and action potential properties in CA1 pyramidal neurons of acute rat hippocampal slices. Front. Synaptic Neurosci. 3, 1–16 (2011).

18. Yang, K. & Dani, J. A. Dopamine d1 and d5 receptors modulate spike timing-dependent plasticity at medial perforant path to dentate granule cell synapses. J. Neurosci. 34, 15888–15897 (2014).

19. Brzosko, Z., Schultz, W. & Paulsen, O. Retroactive modulation of spike timingdependent plasticity by dopamine. Elife 4, 1–13 (2015).

20. Brzosko, Z., Zannone, S., Schultz, W., Clopath, C. & Paulsen, O. Sequential neuromodulation of hebbian plasticity offers mechanism for effective reward-based navigation. Elife 6, 1–18 (2017).

21. Brzosko, Z., Mierau, S. B. & Paulsen, O. Neuromodulation of Spike-Timing-Dependent Plasticity: Past, Present, and Future. Neuron 103, 563–581 (2019).

22. Müller-Dahlhaus, F., Ziemann, U. & Classen, J. Plasticity resembling spike-timing dependent synaptic plasticity: The evidence in human cortex. Front. Synaptic Neurosci. 2, 1–11 (2010).

23. Verhoog, M. B. et al. Mechanisms Underlying the Rules for Associative Plasticity at Adult Human Neocortical Synapses. J. Neurosci. 33, 17197–17208 (2013).

24. Meredith, R. M., Floyer-Lea, A. M. & Paulsen, O. Maturation of Long-Term Potentiation Induction Rules in Rodent Hippocampus: Role of GABAergic Inhibition. J. Neurosci. 23, 11142–11146 (2003).

25. Buchta, W. C., Mahler, S. V., Harlan, B., Aston-Jones, G. S. & Riegel, A. C. Dopamine terminals from the ventral tegmental area gate intrinsic inhibition in the prefrontal cortex. Physiol. Rep. 5, 1–13 (2017).

26. Markram, H., Gerstner, W. & Sjöström, P. J. A history of spike-timing-dependent plasticity. Front. Synaptic Neurosci. 3, 1–24 (2011).

27. Flurkey, K., Currer, J. M. & Harrison, D. Mouse Models of Inherited Human Neurodegenerative Disease. in The mouse in biomedical research. Normative Biology, Husbandry, and Models (eds. Fox, J. G. et al.) vol. 3 809 (Academic Press, 2007).

28. Ferri, S. L. et al. Sociability development in mice with cell-specific deletion of the NMDA receptor NR1 subunit gene. Genes, Brain Behav. 19, 1–12 (2020).

29. Kumar, M. et al. High resolution magnetic resonance imaging for characterization of the Neuroligin-3 knock-in mouse model associated with autism spectrum disorder. PLoS One 9, (2014).

30. Banerjee, A. et al. Double dissociation of spike timing-dependent potentiation and depression by subunit-preferring NMDA receptor antagonists in mouse barrel cortex. Cereb. Cortex 19, 2959–2969 (2009).

31. Sjöström, P. J. & Häusser, M. A Cooperative Switch Determines the Sign of Synaptic Plasticity in Distal Dendrites of Neocortical Pyramidal Neurons. Neuron 51, 227–238 (2006).

32. Letzkus, J. J., Kampa, B. M. & Stuart, G. J. Learning rules for spike timingdependent plasticity depend on dendritic synapse location. J. Neurosci. 26, 10420–10429 (2006).

33. Froemke, R. C., Poo, M. M. & Dan, Y. Spike-timing-dependent synaptic plasticity depends on dendritic location. Nature 434, 221–225 (2005).

34. Kalmbach, B. E. et al. h-Channels Contribute to Divergent Intrinsic Membrane Properties of Supragranular Pyramidal Neurons in Human versus Mouse Cerebral Cortex. Neuron 100, 1194–1208.e5 (2018).

35. Beaulieu-Laroche, L. et al. Enhanced Dendritic Compartmentalization in Human Cortical Neurons. Cell 175, 643–651.e14 (2018).

36. Hodge, R. D. et al. Conserved cell types with divergent features in human versus mouse cortex. Nature 573, 61–68 (2019).

37. Gidon, A. et al. Dendritic action potentials and computation in human layer 2/3 cortical neurons. Science (80-.). 367, 83–87 (2020).

38. Berg, J. et al. Human cortical expansion involves diversification and specialization of supragranular intratelencephalic-projecting neurons. bioRxiv 2020.03.31.018820 (2020) doi: 10.1101/2020.03.31.018820.

39. Kepecs, A. & Fishell, G. Interneuron cell types are fit to function. Nature 505, 318–326 (2014).

40. Bocchio, M. et al. Group II Metabotropic Glutamate Receptors Mediate Presynaptic Inhibition of Excitatory Transmission in Pyramidal Neurons of the Human Cerebral Cortex. Front. Cell. Neurosci. 12, 508 (2019).

41. Komlósi, G. et al. Fluoxetine (Prozac) and serotonin act on excitatory synaptic transmission to suppress single layer 2/3 pyramidal neuron- triggered cell assemblies in the human prefrontal cortex. J. Neurosci. 32, 16369–16378 (2012).

42. Kroon, T. et al. Group I mGluR-mediated activation of martinotti cells inhibits local cortical circuitry in human cortex. Front. Cell. Neurosci. 13, 315 (2019).

43. Palacios-Filardo, J. & Mellor, J. R. Neuromodulation of hippocampal longterm synaptic plasticity. Curr. Opin. Neurobiol. 54, 37–43 (2019).

44. Izhikevich, E. M. Solving the distal reward problem through linkage of STDP and dopamine signaling. Cereb. Cortex 17, 2443–2452 (2007).

45. Sutton, R. S. & Barto, A. G. Introduction to reinforcement learning. vol. 135 (MIT press Cambridge, 1998).

46. Ting, J. T. et al. A robust ex vivo experimental platform for molecular-genetic dissection of adult human neocortical cell types and circuits. Sci. Rep. 8, 1–13 (2018).

47. Ahad, N. A., Yin, T. S., Othman, A. R. & Yaacob, C. R. Sensitivity of normality tests to non-normal data. Sains Malaysiana 40, 637–641 (2011).

