## Supplementary material for "Dopaminergic neuromodulation of spike timing dependent plasticity in mature adult rodent and human cortical neurons": Suppl file

**Supplementary Files**

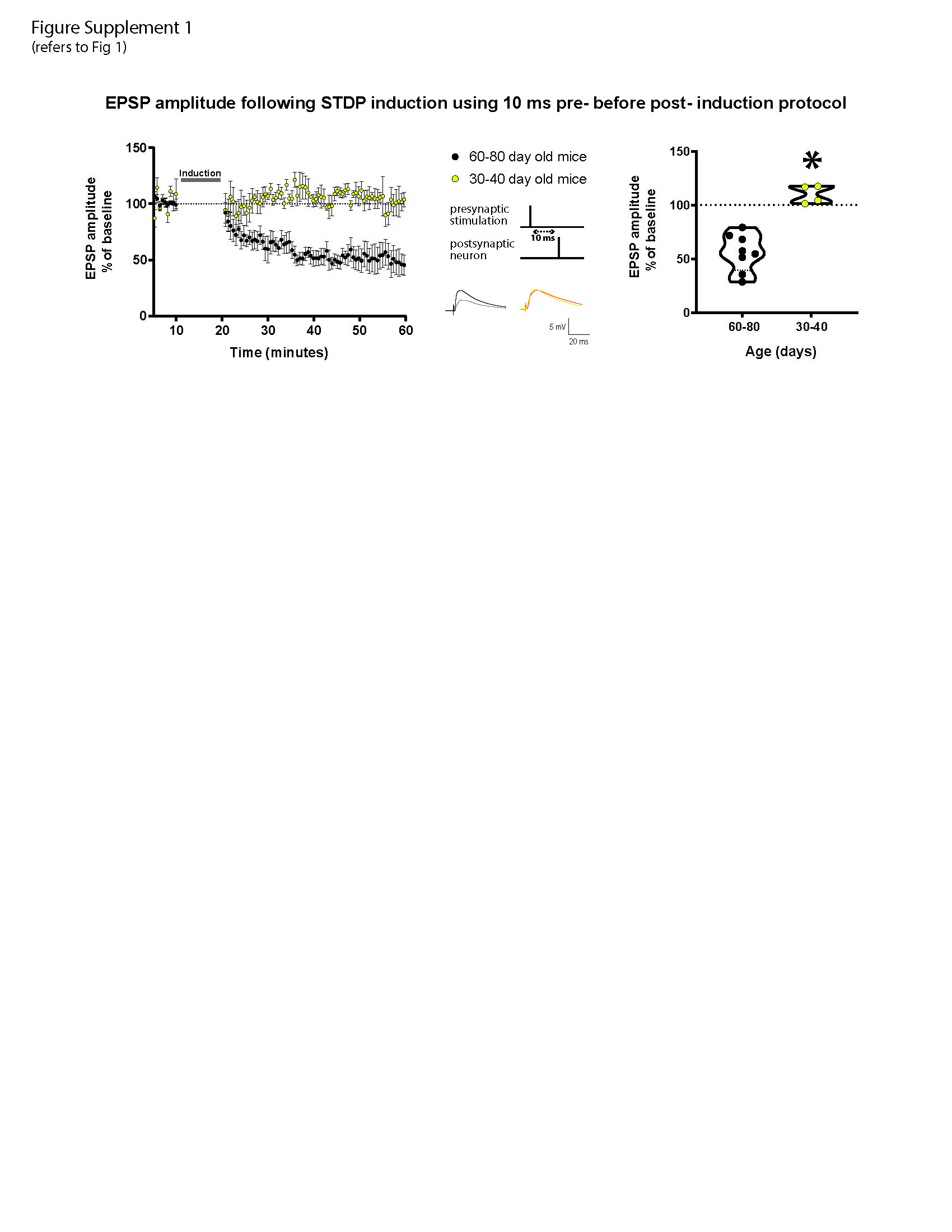

**Figure Supplement 1 (Supplement to Fig 1)**

The peak amplitude of EPSPs following at Δt +10ms STDP induction protocol in mature adult and adult mouse layer 5 cortical pyramidal neurons. Left, the time-course of the EPSP peak amplitude during the STDP experiment in both groups of mice. Middle, the STDP induction protocol timing is illustrated and below are example traces of EPSPs. The baseline trace is the darker trace, the trace following STDP induction is the lighter trace; each trace is the average of 80 traces from the same recording. Right, violin plots of summary data showing that mature adult mice exhibited t-LTD (53.4 ± 8.1, *p* = 0.0002 vs 100 %) whereas adult mice exhibited no change (110.2 ± 4.0 ± 2.9, *p* = 0.1 vs 100 %). There was a significant difference between EPSP peak amplitude following STDP induction in mature adult and adult mice (*p* = 0.0002). All data are shown as mean ± SEM. Number of neurons recorded in mature adult mice, n= 8, and adult mice n= 4.

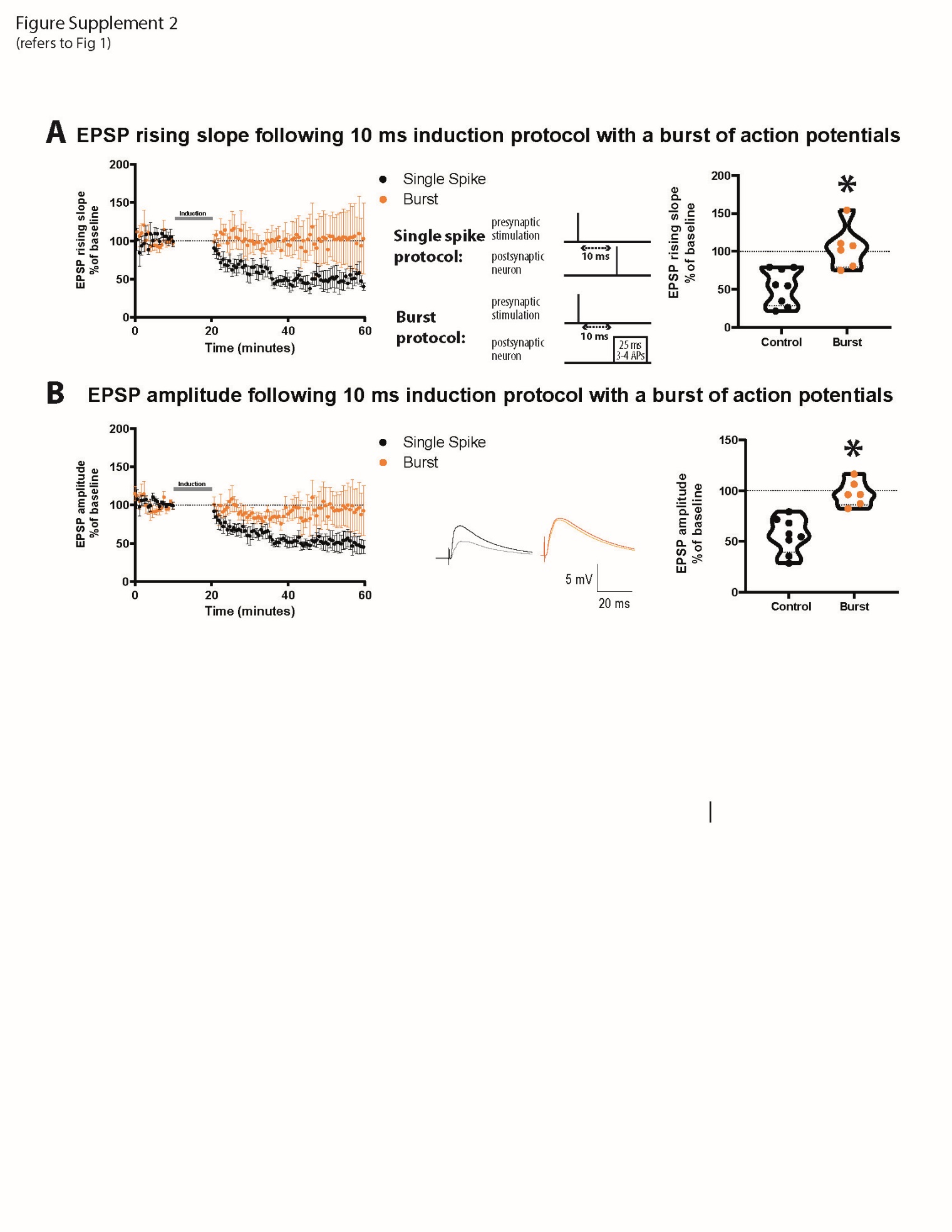

**Figure Supplement 2 (Supplement to Fig 1)**

A burst of APs STDP induction protocol blocks t-LTD in mature adult mice. (A) The time-course of the EPSP rising slope during the STDP experiment is shown to the left. The STDP induction protocol timing is illustrated in the middle. Violin plots of summary of the data showing a significant difference between EPSP rising slope using a single spike or burst protocol (*p* = 0.003) are shown to the right. Using a burst protocol alone results in no change in EPSP rising slope (104.9 ± 11.5, *p* = 0.5 vs 100 %). (B) The time-course of the EPSP amplitude during the STDP experiment. Example traces of EPSPs are shown in the middle. The baseline trace is the darker trace, the resultant trace following STDP induction is the lighter trace; each trace is the average of 80 traces from the same recording. Violin plots of summary of the data showing a significant difference between EPSP amplitude using a single spike or burst protocol (*p* = 0.0003) are shown on the right. Using a burst protocol results in no change in EPSP amplitude (97.4 ± 5.1, *p* = 0.6 vs 100 %). All data are shown as mean ± SEM. For the single spike protocol n = 8 and for the burst protocol n = 6.

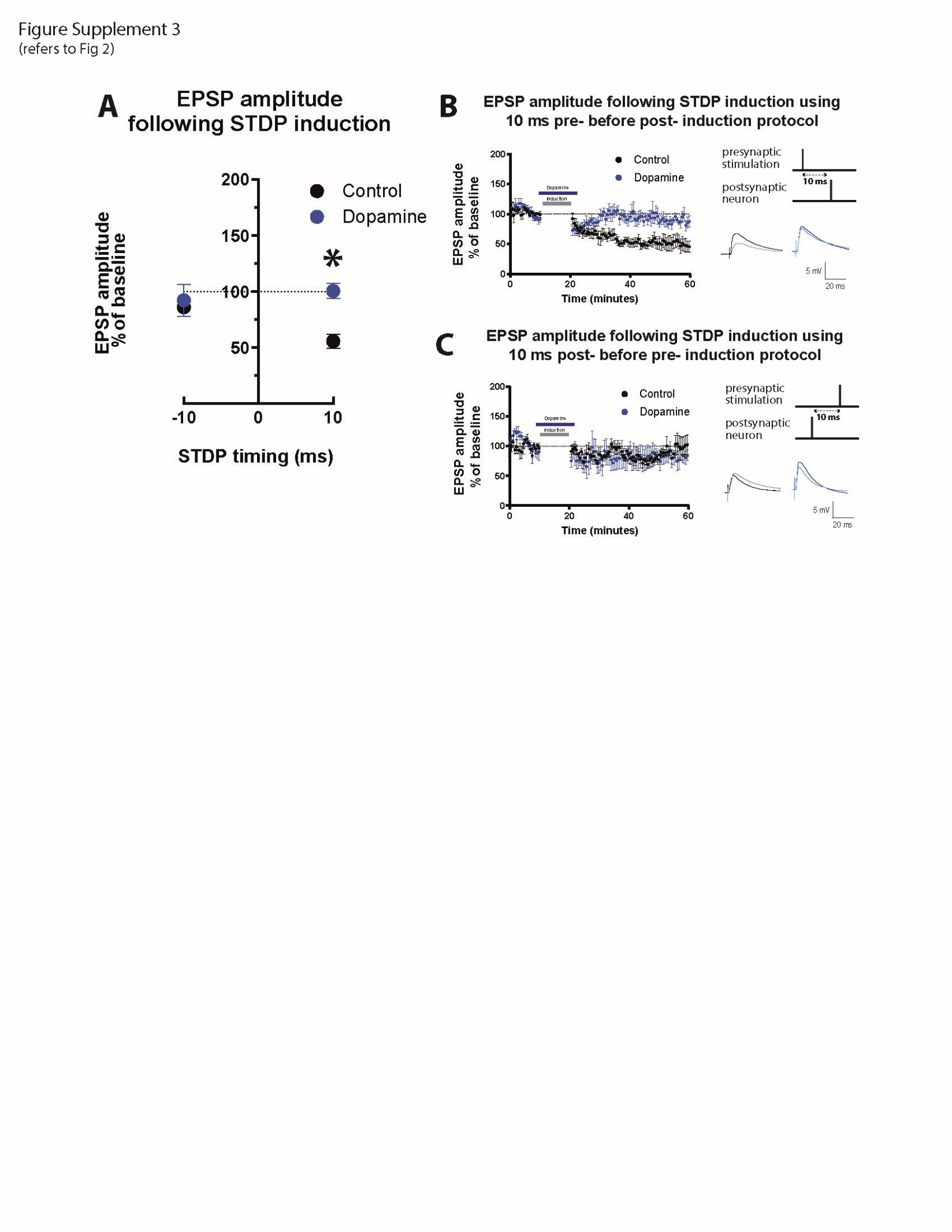

**Figure Supplement 3 (Supplement to Fig 2)**

Effect of DA on EPSP amplitude following STDP induction in mature adult mouse cortical layer 5 pyramidal neurons. (A) STDP induction at Δt -10 ms and + 10 ms EPSP-AP pairing timings with and without 20 μM DA application. EPSP amplitude remained unchanged with the -10ms timing for both the control (88.5 ± 4.7, *p* = 0.6 vs 100%) and DA groups (92.2 ± 14.3, *p* = 0.6 vs 100%) and they were not significantly different from each other (*p* = 0.8). At the +10 ms timing, EPSP amplitude was significantly different between the groups (*p* = 0.0003), resulting in t-LTD in the control group (55.7 ± 6.2, p = 0.0002 vs 100%) and no change in the DA group (97.9 ± 9.0, *p* = 0.9 vs 100 %). The time-course of the EPSP amplitude during the STDP experiment using the +10 ms timing is shown in (B) and the -10 ms timing in (C). DA bath application and time of STDP induction are indicated by bars in the graph. To the right, the STDP induction protocol timing is illustrated and below are example traces of EPSPs. The baseline trace is the darker trace, the resultant trace following STDP induction is the lighter trace; each trace is the average of 80 traces from the same recording. All data are shown as mean ± SEM. For the -10ms timing n = 6 and for the +10ms timing n=8.

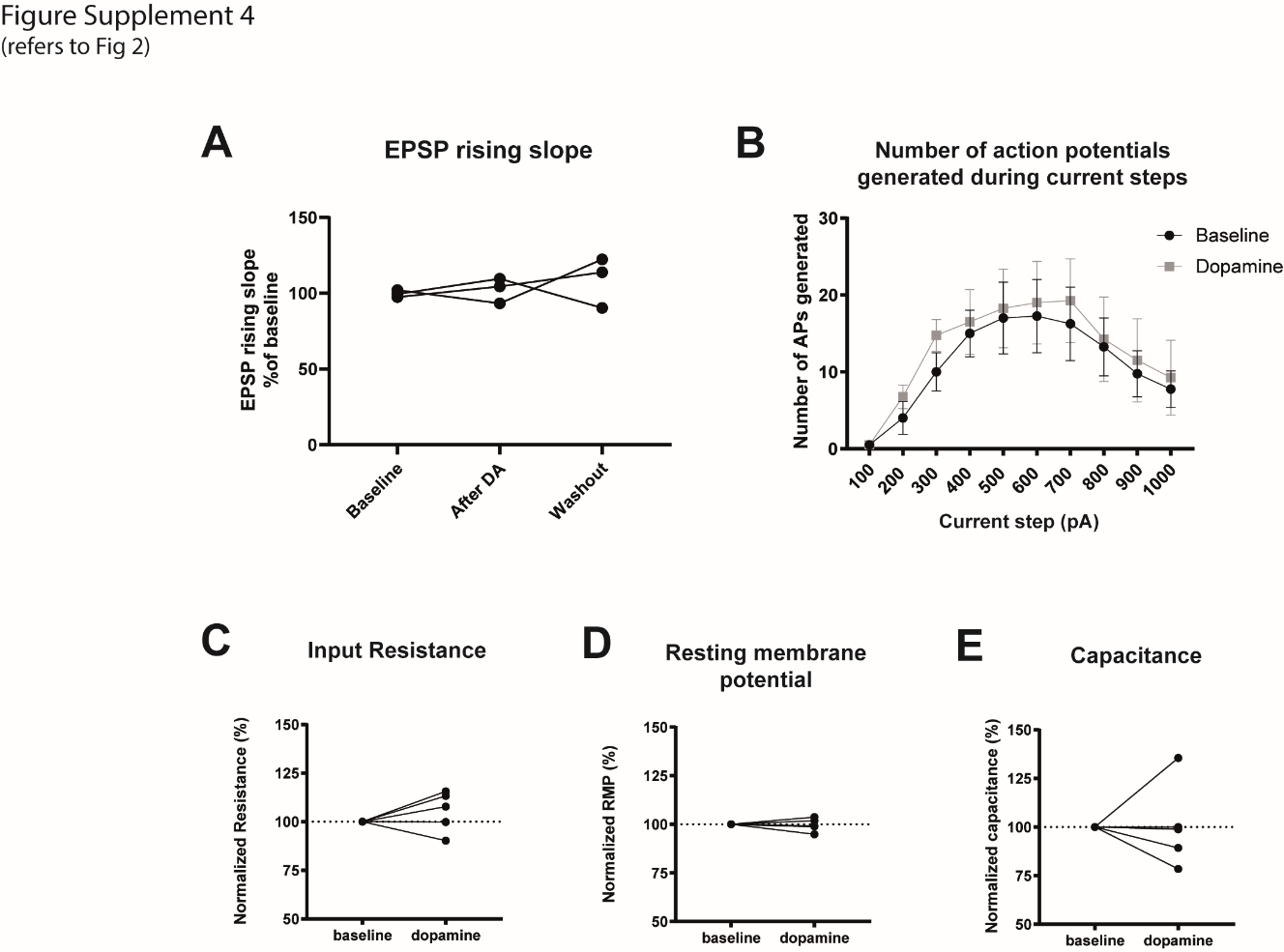

**Figure Supplement 4 (Supplement to Fig 2)**

DA does not affect low frequency-evoked EPSP or basic electrophysiological properties. (A) While evoking EPSPs at 0.14 Hz, the same rate as baseline stimulation before STDP induction, 20 μM DA was bath applied without STDP induction for seven minutes. Data points were taken one minute before DA application (control), during the final minute of DA application (DA data) and ten minutes after the end of DA application (washout). DA application had no effect on EPSP rising slope (repeated measures one-way ANOVA, *p* = 0.6 F_(1.1, 2.2)_ = 0.4). We also measured effect of DA on AP firing frequency (B, two-way ANOVA, effect of DA: *p* = 0.3 F_(1, 60)_ = 1.2, effect of current step: *p* = 0.0002 F_(9, 60)_ = 4.5, interaction: *p* = 0.9 F_(9, 60)_ = 0.05), input resistance (C, DA: 105.4 ± 4.6, *p* = 0.3 vs 100%), resting membrane potential (D, DA: 99.6 ± 1.5, *p* = 0.8 vs 100%) and capacitance (E, DA: 100.4 ± 9.6, *p* = 0.9 vs 100%). DA did not significantly affect these parameters. Individual data points are shown, except in B where data is shown as mean ± SEM. For EPSP rising slope, n = 3. For the remaining n = 5.

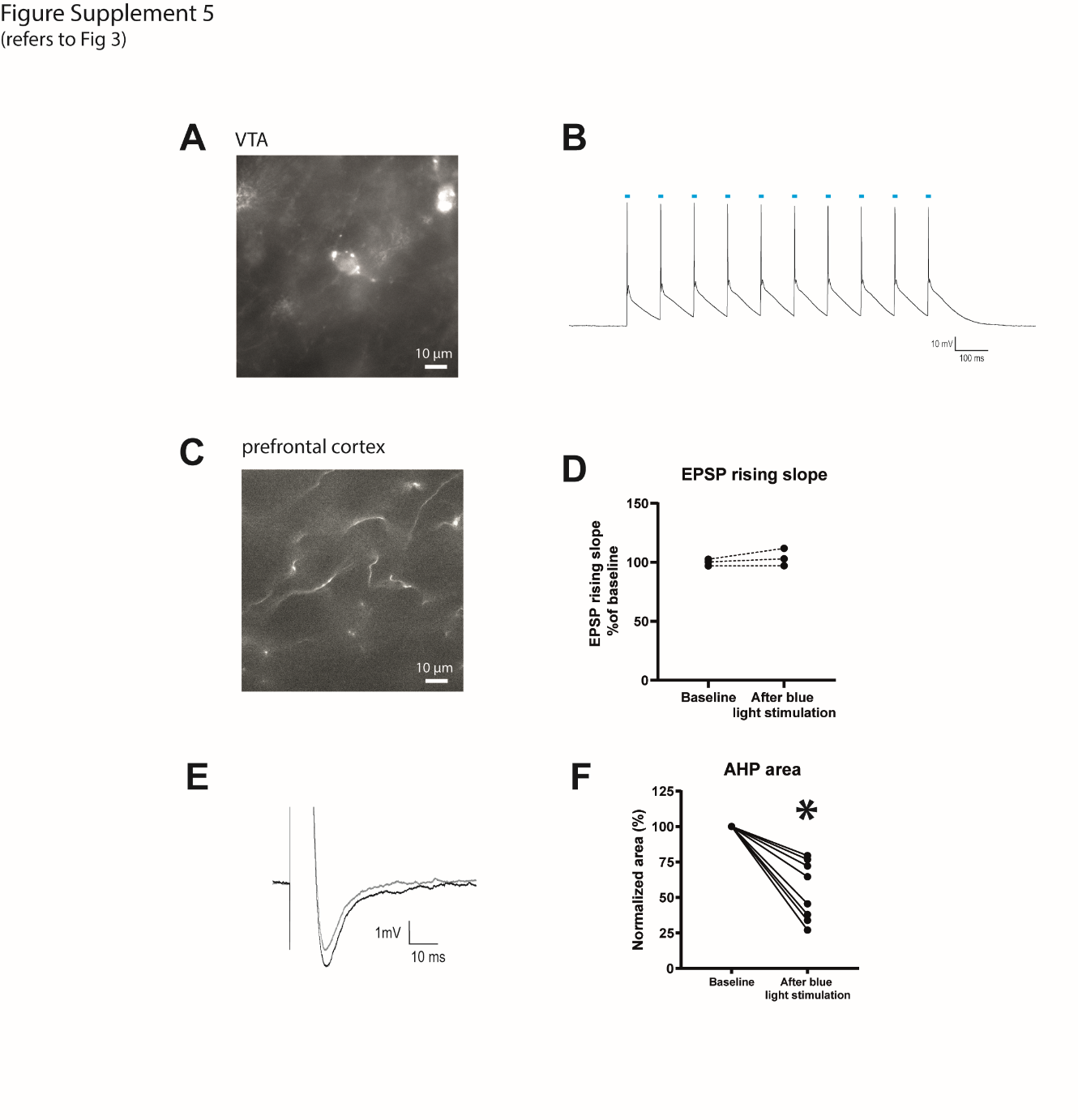

**Figure Supplement 5 (Supplement to Fig 3)**

Viral transfection of ChR2 in dopaminergic neurons of the VTA from Dat^IREScre^ mice. (A) An example of a VTA cell with viral expression as demonstrated by eYFP fluorescence. (B) Optogenetic stimulation of these cells with a 10 Hz pulse train of blue light pulses (shown as blue bars) evoked APs in current clamp mode. (C) example of prefrontal cortex fibers from VTA neurons as demonstrated by eYFP fluorescence. (D) Blue light stimulation alone had no effect on EPSP rising slope (*p* = 0.3, n = 3). (E) Averaged AHP traces before and after blue light stimulation. The baseline trace is the darker trace, the after trace is the lighter trace; each trace is the average of 3 traces from the same recording. (F) AHP area analysis showing that AHP area is decreased following blue light stimulation (*p* = 0.0005, n = 8).

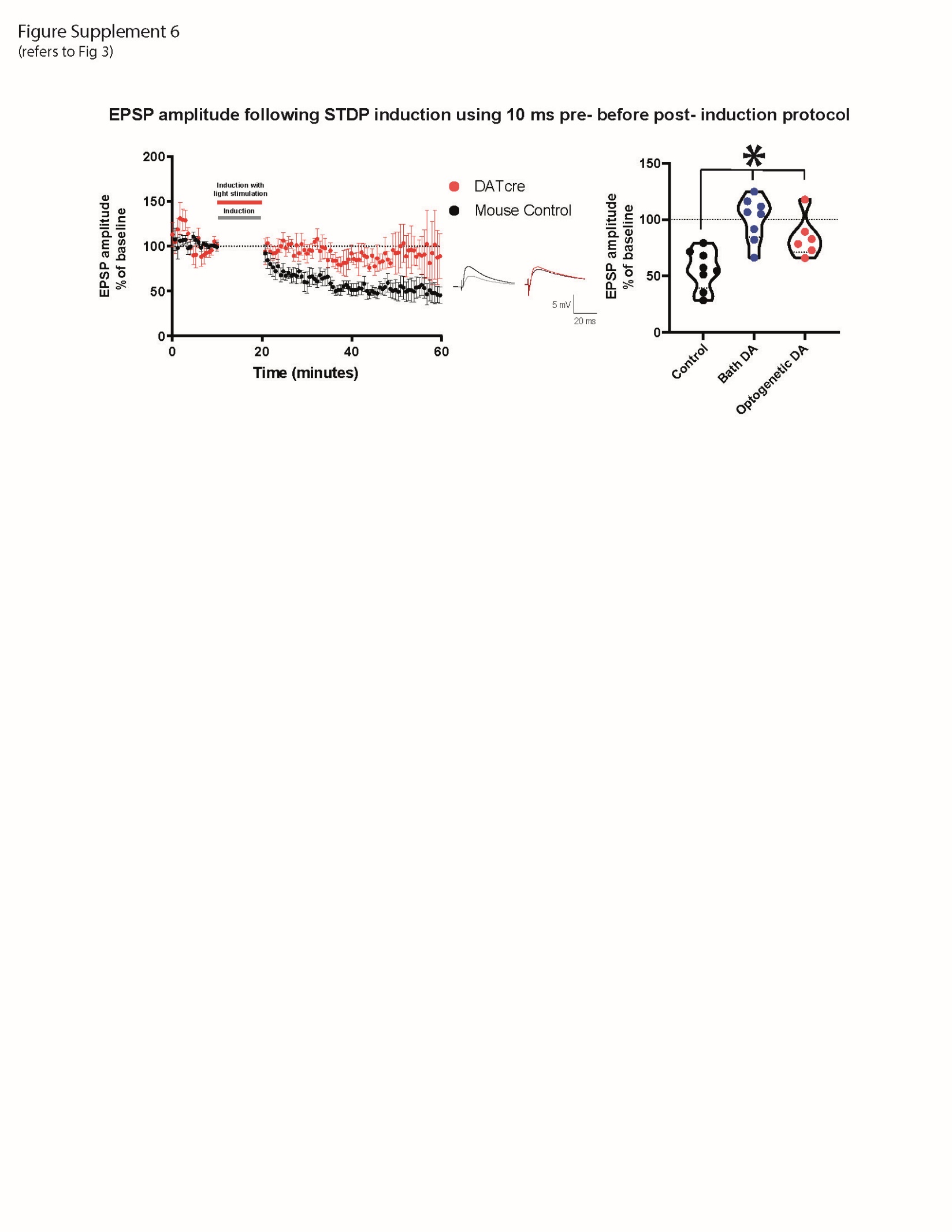

**Figure Supplement 6 (Supplement to Fig 3)**

Optogenetically triggered DA release during STDP induction blocks EPSP peak amplitude t-LTD in cortical layer 5 pyramidal neurons of mature adult mouse. The time-course of the EPSP peak amplitude during the STDP experiment in both control and Dat^IREScre^ mice (left) and violin plots of summary of the data (right). Example EPSP traces are shown (middle). The baseline trace is the darker trace, the after trace is the lighter trace; each trace is the average of 80 traces from the same recording. Data show that, similar to bath application of DA, optogenetically triggered release of DA shows no change in EPSP amplitude (85.2 ± 7.2, *p* = 0.1 vs 100 %, paired *t*-test). A significant difference between the control group and the DA exposed groups was detected (one way ANOVA, *p* = 0.0004, F = 12.2_(2, 19)_ , Tukey’s multiple comparison test, control vs bath application of dopamine: *p* = 0.0003, control vs optogenetic DA: *p* = 0.02, bath vs optogenetic application of DA: *p* = 0.3). All data are shown as mean ± SEM. For the optogenetic stimulation n = 6, while for the control and bath DA groups n = 8.

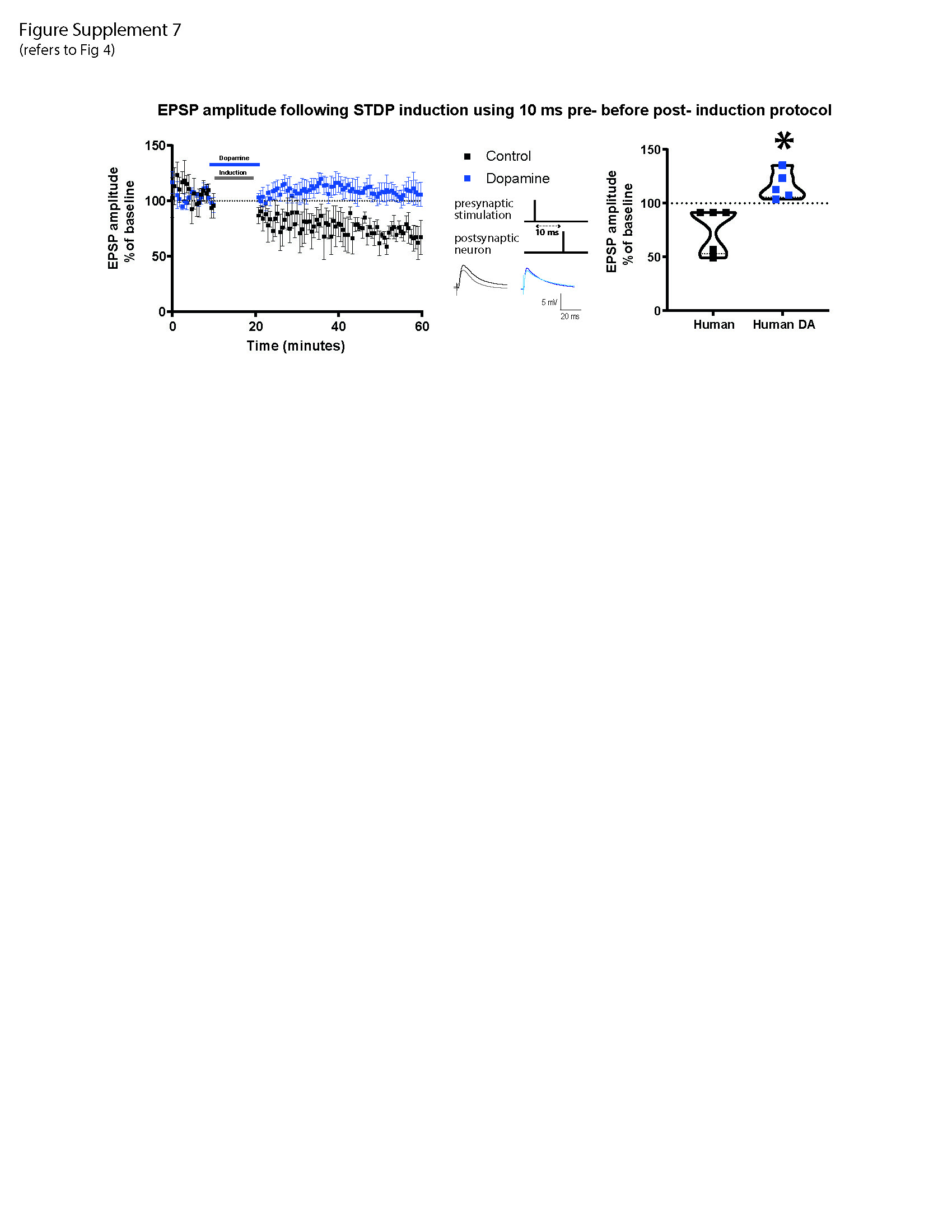

**Figure Supplement 7 (Supplement to Fig 4)**

DA potentiates baseline EPSP peak amplitude after Δt +10ms STDP protocol in adult human cortical layer 5 pyramidal neurons. Left, the time-course of the EPSP amplitude after Δt +10ms STDP induction protocol. Middle, the STDP induction protocol timing is illustrated and below are example traces of EPSPs. The baseline trace is the darker trace, the resultant trace following STDP induction is the lighter trace; each trace is the average of 80 traces from the same recording. Right, violin plots of summary of the results showing a trend toward t-LTD in the control group (76.1 ± 9.5, *p* = 0.07 vs 100 %) and a trend toward t-LTP in the DA group (116.6 ± 5.8, *p* = 0.05 vs 100 %). There was a significant difference between EPSP amplitude with and without DA application during STDP induction (*p* = 0.007). All data are shown as mean ± SEM. For both groups, n = 5.

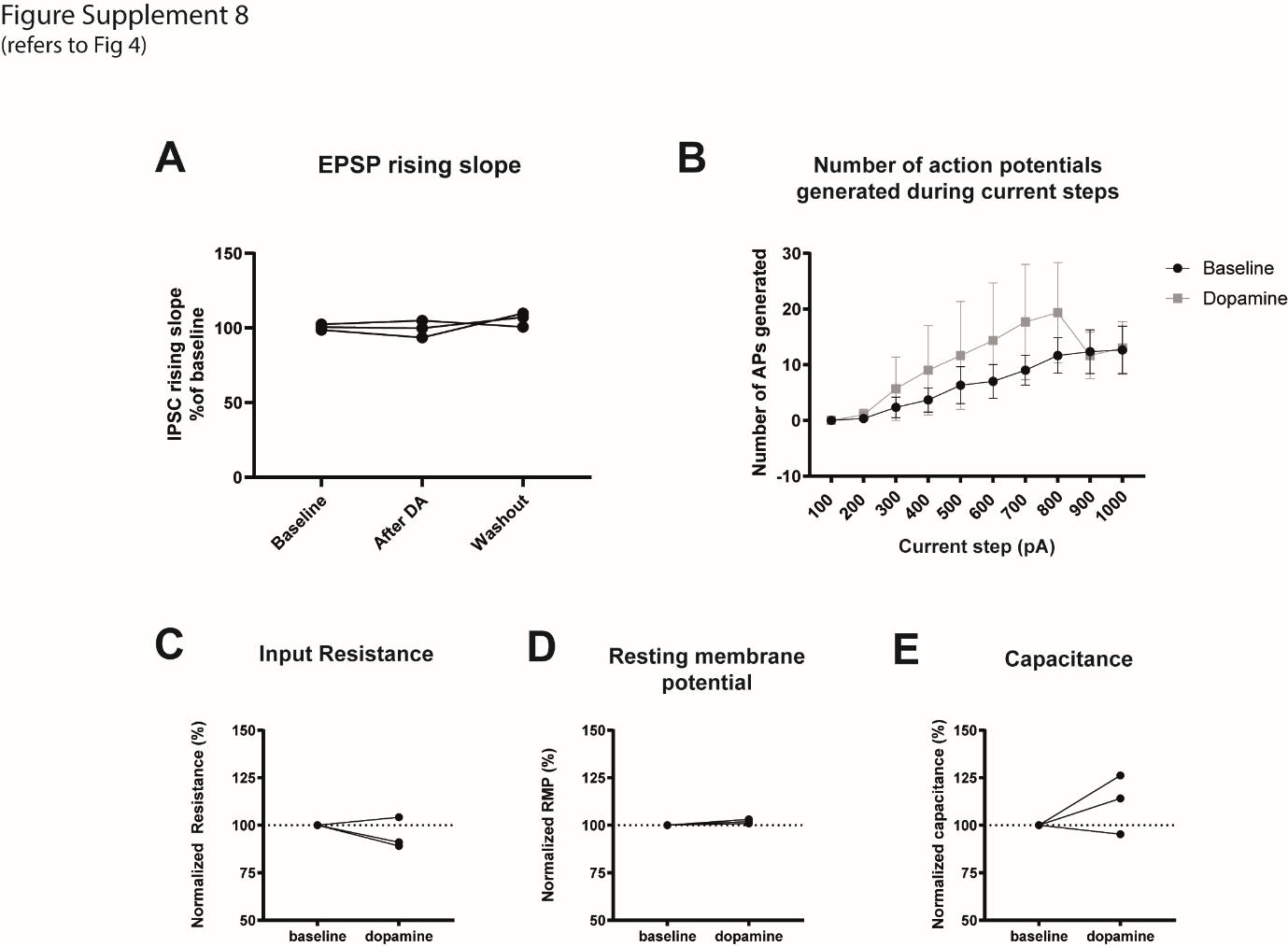

**Figure Supplement 8 (Supplement to Fig 4)**

DA does not affect low frequency-evoked EPSP or basic electrophysiological properties recorded from layer 5 pyramidal neurons of human cortex. (A) EPSPs were evoked at 0.14 Hz, the same rate as the baseline protocol; DA was bath applied without STDP induction for seven minutes. Data points were taken one minute before DA application (control), during the final minute of DA application (DA data) and ten minutes after the end of DA application (washout). DA had no effect on EPSP rising slope (repeated measures one-way ANOVA, *p* = 0.4 F_(1, 2)_ = 1.3). We also measured the action of DA on AP firing frequency (B, two-way ANOVA, effect of dopamine: *p* = 0.1 F_(1, 40)_ = 2.4, effect of current step: *p* = 0.07 F_(9, 40)_ = 2.0, interaction: *p* = 0.9 F_(9, 40)_ = 0.2), input resistance (C, dopamine: 94.8 ± 4.7, *p* = 0.4 vs 100%), resting membrane potential (D, dopamine: 101.9 ± 0.7, *p* = 0.1 vs 100%) and capacitance (E, dopamine: 111.9 ± 9.0, *p* = 0.3 vs 100%). DA did not significantly affect these parameters. Individual data points are shown, except in B where data is shown as mean ± SEM. For all groups, n=3.

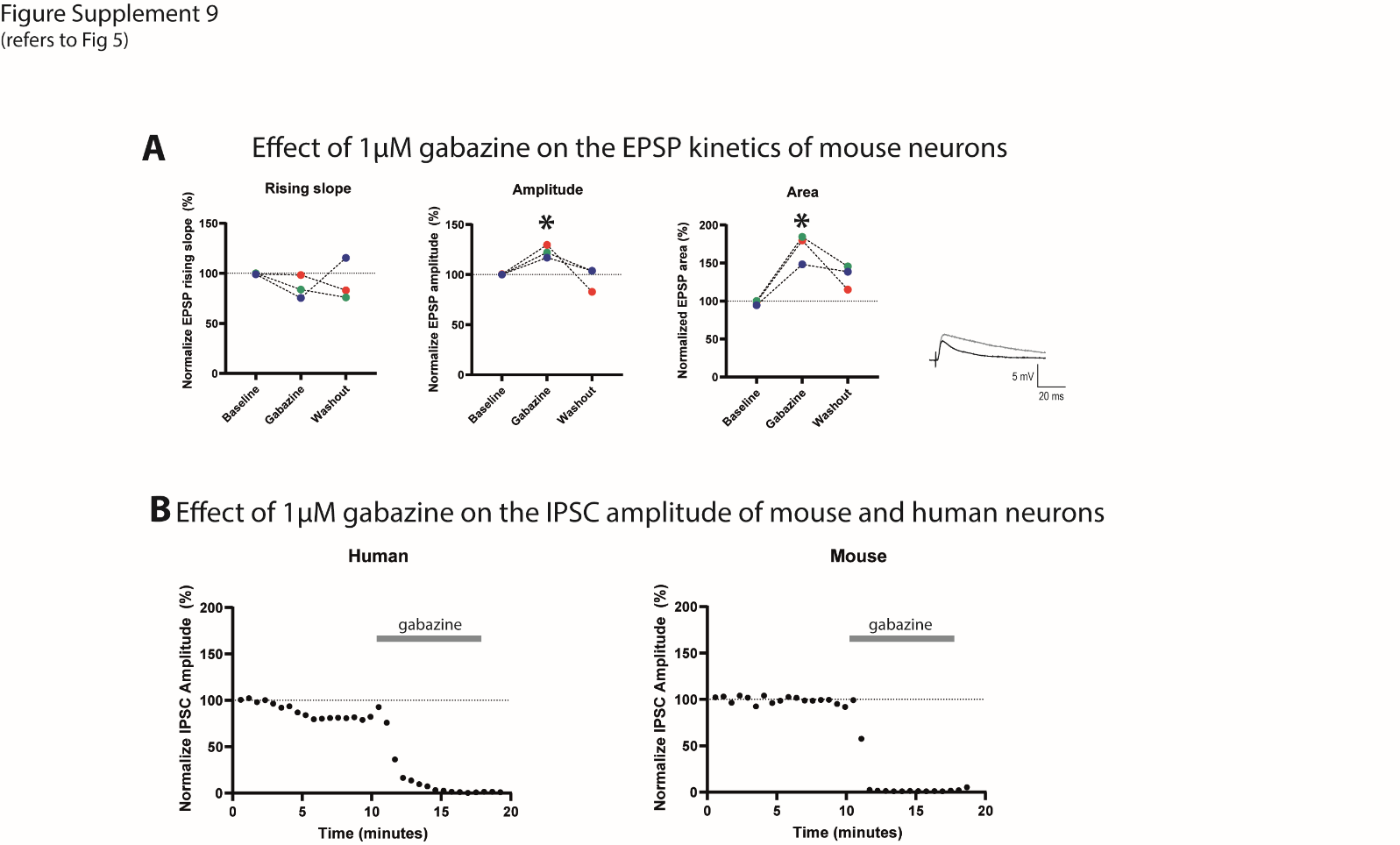

**Figure Supplement 9 (Supplement to Fig 5)**

The GABA-A receptor antagonist gabazine (1 µM) affects EPSCs and abolished IPSCs in human and mouse cortical layer 5 pyramidal neurons. (A) Gabazine application, enhanced EPSP peak amplitude (middle, repeated measures one-way ANOVA, *p* = 0.1, F_(1, 2)_ = 7.0, Dunnett’s multiple comparison baseline vs gabazine *p* = 0.04, baseline vs washout *p* = 0.9) and area (right, repeated measures one-way ANOVA, *p* = 0.1, F_(1, 2)_ = 5.5, Dunnett’s multiple comparison baseline vs gabazine *p* = 0.03, baseline vs washout *p* = 0.1), and did not affect rising slope (left, repeated measures one-way ANOVA, *p* = 0.5, F_(1.1, 2.2)_ = 0.6, Dunnett’s multiple comparison baseline vs gabazine *p* = 0.3, baseline vs washout *p* = 0.8). Individual data points are shown (n = 3), example traces are shown on the far right. The baseline trace is the darker trace, the resultant trace following gabazine application is the lighter trace; each trace is the average of five traces. (B) Recorded IPSCs are abolished by gabazine application (denoted by grey bars). Individual example traces in human (left) and mouse (right) are shown.

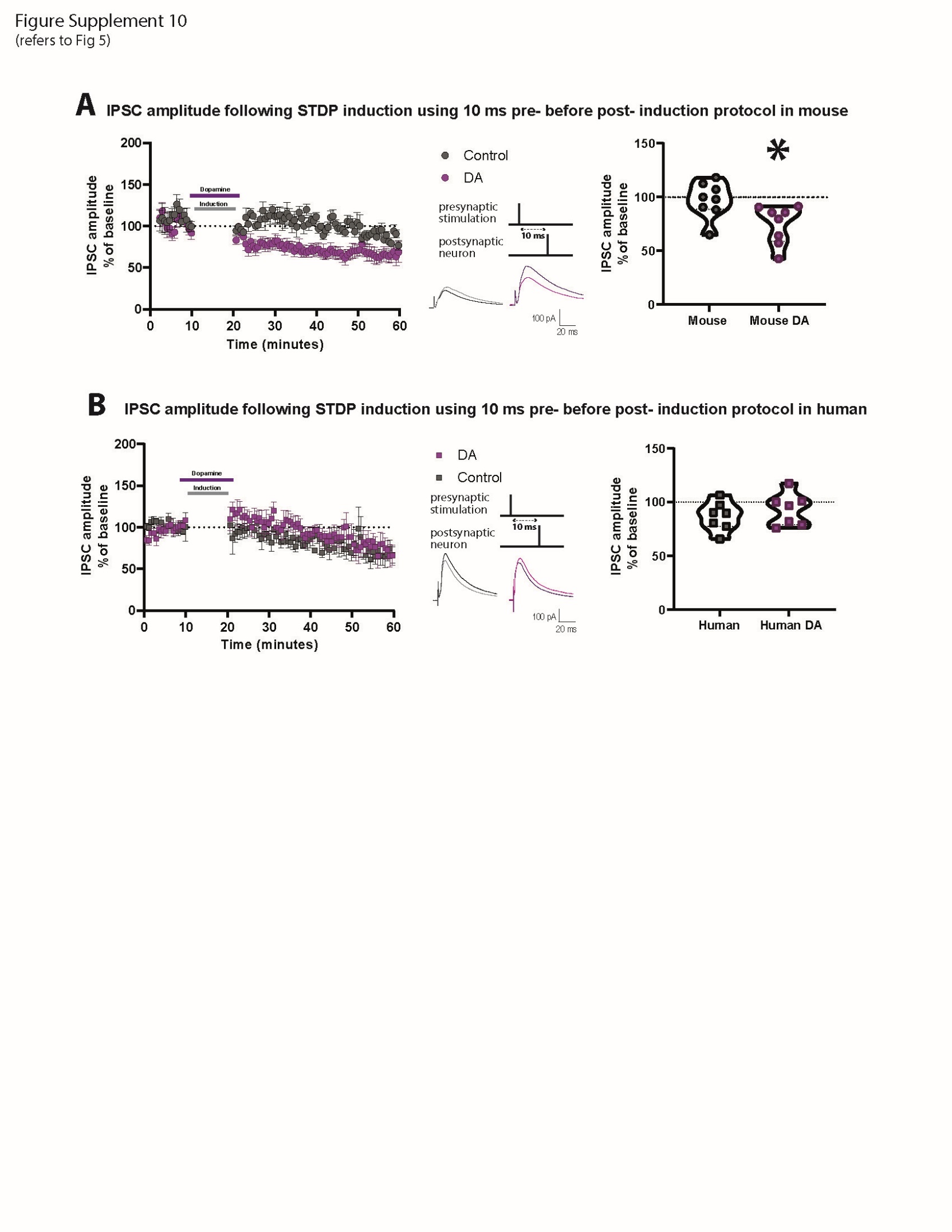

**Figure Supplement 10 (Supplement to Fig 5)**

DA depresses IPSC amplitude after Δt +10ms STDP protocol in cortical layer 5 pyramidal neurons of mature adult mice but not humans. Effect of DA on IPSC amplitude following STDP induction in adult mouse (A; control: 97.5 ± 5.9, *p* = 0.7 vs 100%; DA: 74.7 ± 6.3, *p* = 0.005 vs 100%) and human (B; control: 87.2 ± 5.3, *p* = 0.05 vs 100%; DA: 93.1 ± 5.7, *p* = 0.3 vs 100%) pyramidal neurons. Left, the time-course of the IPSC amplitude after Δt +10ms STDP timing protocol. Middle, the STDP induction protocol timing is illustrated and below are example traces of IPSCs. The baseline trace is the darker trace, the resultant trace following STDP induction is the lighter trace; each trace is the average of 80 traces from the same recording. Right, violin plots showing summary of the results. DA significantly reduced the IPSC amplitude after STDP induction in neurons recorded from mature adult mice (*p* = 0.02, n = 8), but not from humans (*p* = 0.5, n = 7). All data are shown as mean ± SEM.

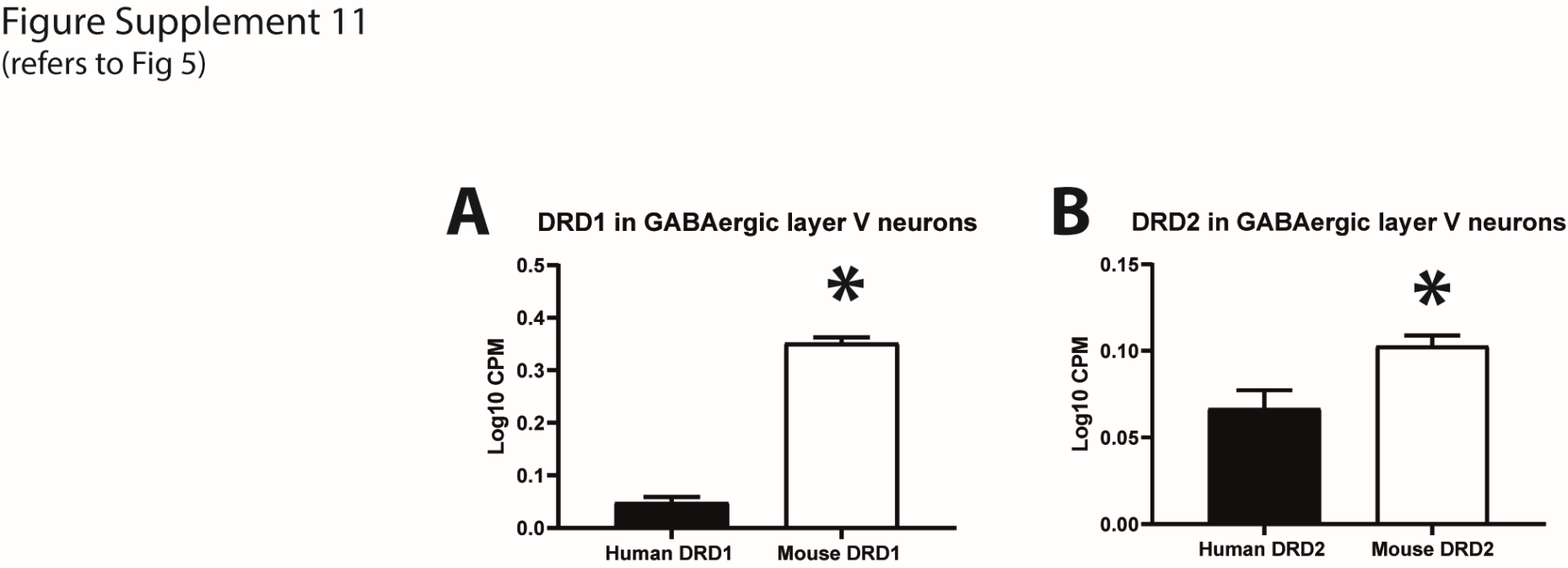

**Figure Supplement 11 (Supplement to Fig 5)**

DA receptor expression data in cortical layer 5 neurons from the Allen Brain Atlas of RNA-seq data. (A) DRD1 expression in cortical layer 5 GABAergic neurons from human (0.05 ± 0.01, n = 786) and mouse (0.4 ± 0.01, n = 3504) are significantly different (*p* = 0.0001). (B) DRD2 expression in cortical layer 5 GABAergic neurons from human (0.07 ± 0.01, n = 786) and mouse (0.1 ± 0.006, n = 3504) are significantly different (*p* = 0.003). Data were not normally distributed and therefore analyzed with Welch’s *t* test.

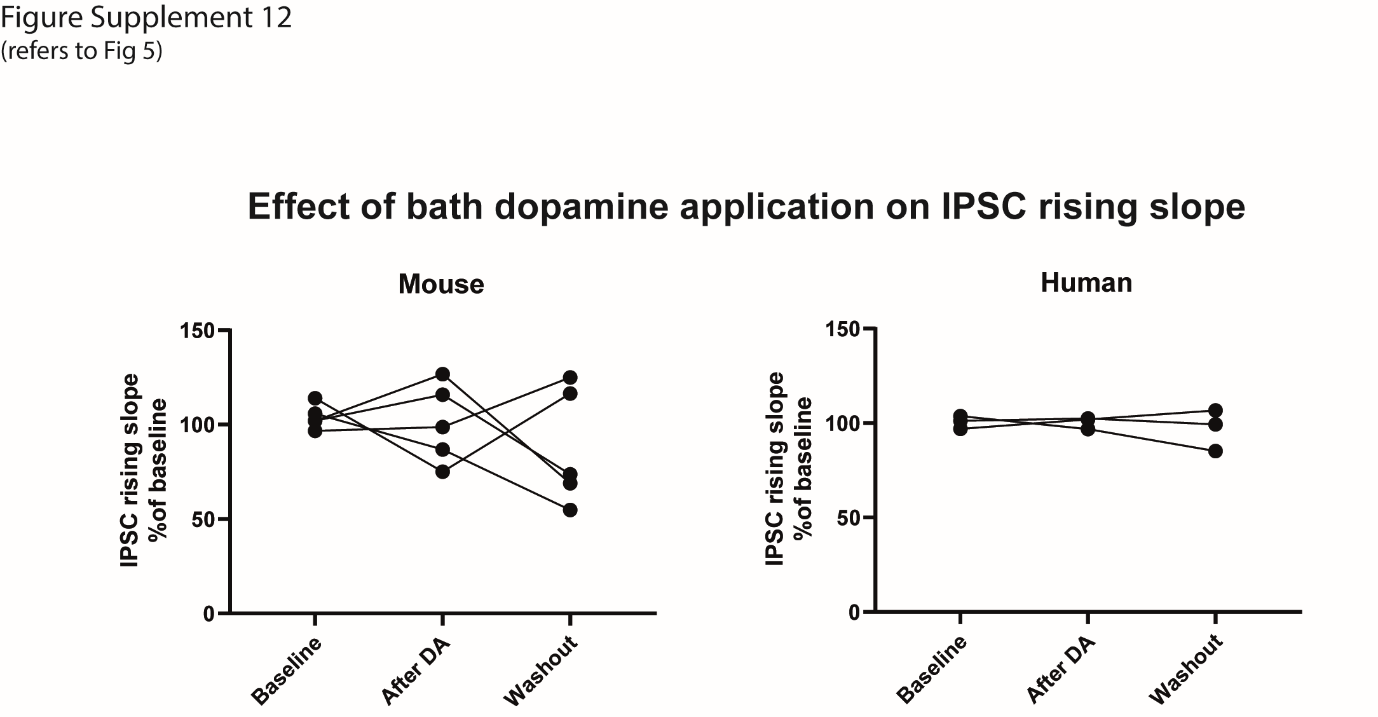

**Figure Supplement 12 (Supplement to Fig 5)**

Action of DA on IPSC rising slope in mouse and human pyramidal neurons. DA was applied for seven minutes while evoking IPSCs at 0.14 Hz, that was the same repetition rate used to collect baseline IPSCs before the STDP protocol. Data points were taken one minute before DA application (control), during the final minute of DA application (DA data) and ten minutes after the end of DA application (washout). DA had no effect on IPSC rising slope in mouse (repeated measures one-way ANOVA, *p* = 0.5, F_(1.4, 5.7)_ = 0.6, n = 5) and human (repeated measures one-way ANOVA, *p* = 0.7, F_(1, 2)_ = 0.2, n = 3) neurons. Individual data points are shown.
